# Symbiotic solutions for colony nutrition: conserved nitrogen recycling within the bacterial pouch of *Tetraponera* ants

**DOI:** 10.1101/2025.03.13.643113

**Authors:** Mingjie Ma, Biru Zhu, Dayong Zhang, Piotr Łukasik, Yi Hu

## Abstract

Microbial symbioses are fundamental to the nutrition of many animal groups, yet the mechanisms of nutrient provisioning remain poorly understood — even in social insects, a diverse clade whose global ecological dominance and sophisticated caste systems make them disproportionately well-studied models for symbiosis research. Here, we investigate the functional significance of a specialized and unusual symbiotic organ— the bacterial pouch—in four ant species within the *Tetraponera nigra-*group, focusing on the symbionts’ roles in nitrogen recycling and colony fitness. Bacterial pouch houses a microbial community consistently dominated by the core bacterial symbiont *Tokpelaia*. Metagenomic sequencing and targeted ¹⁵N-labeled urea feeding experiments demonstrate that these symbionts assimilate nitrogen from urea into amino acids, transferring them to adult workers and developing larvae that incorporate recycled nitrogen into their tissues. Disruption of this nitrogen-recycling symbiosis severely impairs larval growth and overall colony fitness. Our study highlights the critical role of the bacterial pouch in sustaining colony fitness in nitrogen-limited environments, providing new insights into the pivotal role of microbial symbioses in the ecological success of social insects.

## Introduction

Symbiotic relationships have significantly influenced the ecological dynamics and evolutionary trajectories of multicellular organisms as well as their associated microbes. Among the most impactful are symbioses with microorganisms that inhabit dedicated host organs. Classic examples include plant root nodules, which harbor nitrogen-fixing rhizobia^1,2^, and the light organs of bobtail squid^3,4^, home to bioluminescent bacteria. In insects, the well-known examples include mycangia in ambrosia beetles^5^ serving as reservoirs for fungal symbionts, specialized gut organs in bean bugs and other heteropterans^6^, and bacteriomes in most other sap-feeding Hemipteran insects^7,8^ providing a habitat for nutrient-provisioning bacteria. Despite the importance of symbiont-hosting organs, our understanding of the diversity and distributions of these specialized structures, as well as the specificity and ecological significance of symbiotic microbes they harbor, remains notably incomplete.

Ants (Hymenoptera: Formicidae) are among the most ecologically successful and significant groups of insects on Earth. Their mutualistic interactions with microbes have been identified as an important factor driving their long-term evolution and ecological dominance^9,10^. In particular, microbes’ abilities to provide essential nutrients such as amino acids, vitamins, and other metabolic byproducts that ants cannot synthesize themselves are considered a key driver that enables ants to thrive in various nutrient-imbalanced niches, contributing to their widespread ecological dominance^11–15^.

A remarkable pattern in ant evolution is the number of independent shifts from nitrogen-rich carnivorous and omnivorous diets to nitrogen-poor herbivory. Herbivorous ants typically scavenge for foods such as honeydew from tended sap-feeding insects, extrafloral nectar, pollen, fungi, and plant wound secretions^16–18^. Accessible nitrogen and essential amino acids in these food sources are typically limited. The relatively high occurrence of specialized bacteria in herbivorous ant taxa indicates that they may often rely on microbial partners to mitigate nitrogen limitation and nutritional imbalance^9,11,19,20^. Recent studies using high-throughput sequencing and functional assays have shown a trend of functional convergence in nitrogen recycling capabilities among symbionts of unrelated herbivorous ants that feed on N-poor or N-inaccessible foods. For example, in carpenter ants (genus *Camponotus*), intracellular symbiont *Blochmannia* synthesizes essential amino acids from recycled nitrogen for their hosts^21,22^. In the herbivorous ant tribe Cephalotini, analogous roles fulfilled by abundant, conserved multi-species bacterial communities present within the gut lumen were both predicted from metagenomic data and demonstrated *in vivo* and *in vitro*^15,23^.

Social insects like ants face heightened nutrient regulation complexity compared to solitary species due to division of labor and intra-colony variability. Only a subset of workers (foragers) gather food, requiring them to balance their own nutritional needs with the diverse demands of the entire colony^24^. Larvae require substantial protein intake for their rapid growth, and adult workers mostly need carbohydrates for energy^24^. This dietary divergence presents a greater challenge for ant larvae in obtaining sufficient nitrogen when compared to adult workers. The critical role of symbiotic bacteria in colony development through their impact on nitrogen balance and brood nutrition has been demonstrated in *Camponotus* ants, which harbor intracellular symbionts *Blochmannia* at all developmental stages. It was shown that antibiotic-treated colonies suffered from a decline in the number of larvae developing into pupae, smaller body size of newly eclosed workers, and reduced overall colony sizes^25^. In contrast, extracellular symbionts associated with ants are typically restricted to specific life stages, largely due to the drastic tissue remodeling during the metamorphosis^26^. A critical unanswered question is how these life stage-specific extracellular symbionts, if any, could contribute to the nutritional needs of colony members at different life stages in the entire colony.

The arboreal ant genus *Tetraponera,* restricted to the Palaeotropics and Australia, encompasses approximately 110 described species divided into six distinct species groups^27,28^ that were shown to vary in their nutritional biology and anatomy^29–31^. The *T. nigra*-group, comprising 20 species, is uniquely characterized by the presence of an unusual bacterial pouch located at the junction between the midgut and hindgut. This pouch is connected to Malpighian tubules and the tracheal system and is densely populated with bacteria^32^. This structure bears a resemblance to the symbiotic organs observed in certain clades of other insects, including leaf beetles^33^, bean bugs^34^, and ambrosia beetles^5^, suggesting that these ants engage in a specialized form of nutritional symbiosis with microbial partners^35^. Stable isotopic analysis revealed that *T. nigra*-group ants are at a relatively low trophic level (Fig. 1b). Indeed, three species in the group, *T. tucurua*^36^, *T. binghami*^31,37^, and *T. punctulata*^36^, have been documented feeding on honeydew produced by scale insects, and one species, *T. rotula*^38^, feeding on extrafloral nectar —foods considered nutritionally imbalanced. We hypothesized that the nutritional symbiosis with specialized microorganisms housed within gut pouches may have enabled these ants to occupy ecological niches where essential nutrients are otherwise scarce, providing a potential explanation for the success of this group in such habitats. Hence, in a broad sample spanning multiple colonies of four *Tetraponera* species (Supplementary Data 1), we anticipated a stable symbiosis, with microbes benefiting hosts through nitrogen recycling and nutrient biosynthesis.

**Figure 1.**
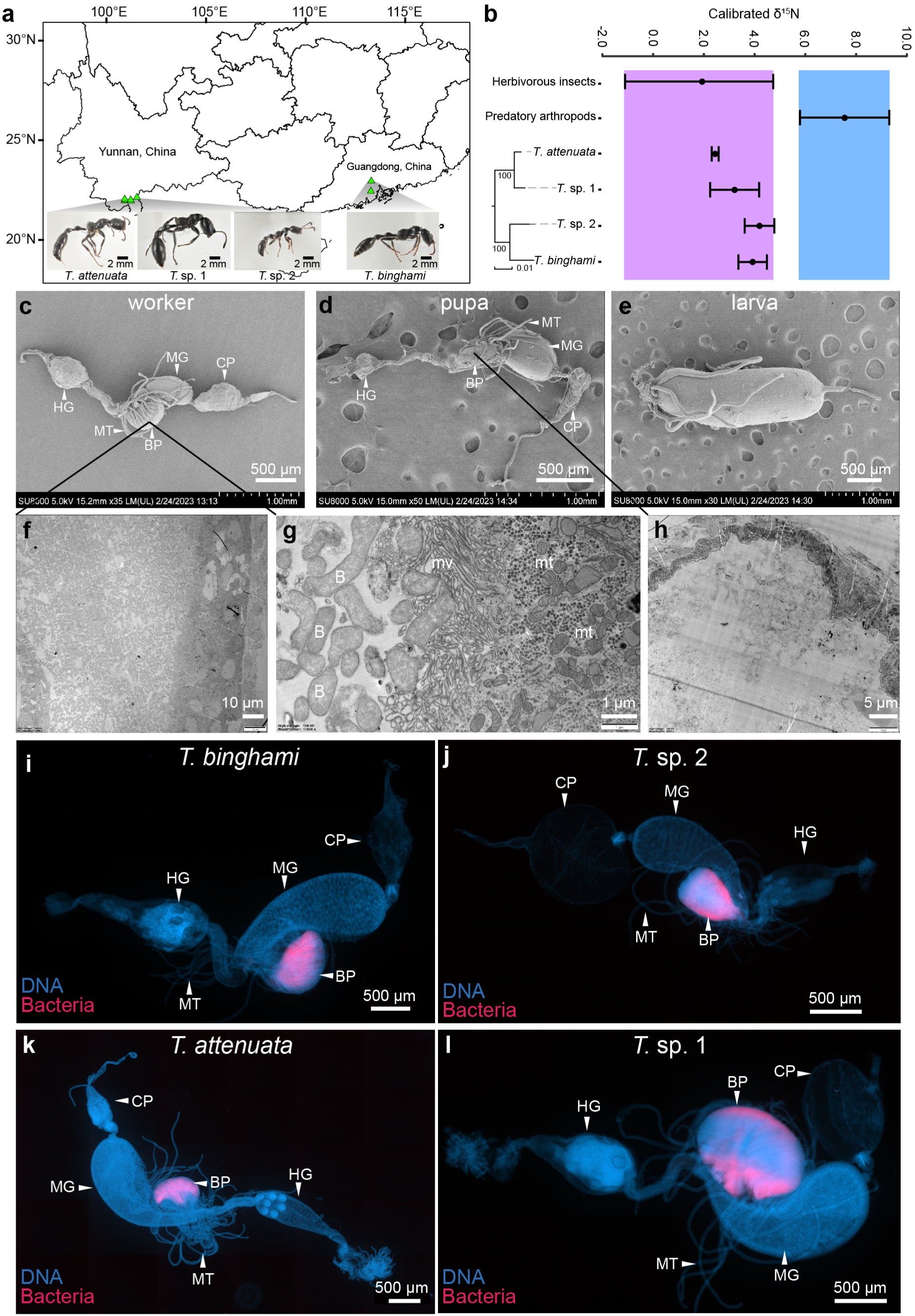
Specimen origins and the gut structure organization in the *T. nigra*-group ants. **(a)** Sampling locales for four ant species used in this study (image credit: Mingjie Ma). **(b)** Nitrogen isotope ratios (mean +/− S.D.) were utilized to assess the trophic levels of the four species through comparisons against herbivorous insects (purple box) and predatory arthropods (blue box) from the same site. The phylogeny of the four ant species is based on 1329 single-copy orthologs. **(c-e)** Scanning electron microscope (SEM) images showing the gut structure of *T. attenuata* at different developmental stages: worker, pupa, and larva. **(f-h)** Transmission electron microscope (TEM) images of ultrathin sections illustrating the bacteria within the bacterial pouch of *T. attenuata* at worker (with different magnifications) and pupal stage, respectively. **(i-l)** Fluorescence in situ hybridization (FISH) of bacteria within the bacterial pouch of four *Tetraponera* ant species. DNA (DAPI) and broad-spectrum bacterial probe signals are denoted by blue and magenta, respectively. CP, crop; MG, midgut; MT, Malpighian tubules; BP, bacterial pouch; HG, hindgut; B, bacteria; mv, microvillar differentiation; mt, mitochondria.

The goal of this study was to comprehensively characterize the diversity and functional roles of bacterial symbionts and their significance in the nutritional biology of the *T. nigra-*group ants. We first explored the general organization of the gut structure in connection with the bacterial pouch across ant life stages. Second, we characterized the diversity of microbial communities within bacterial pouches across colonies of four species and the phylogenetic position and genomic characteristics of the dominant bacteria. Third, we assessed the roles of these symbiotic bacteria in nutrient metabolism, with an emphasis on nitrogen recycling and provisioning. Lastly, we explored how symbionts contribute to nutritional supplementation for the entire colony, particularly for larvae. We did this by combining microscopic analysis, 16S rRNA gene amplicon sequencing, metagenomics, and *in vivo* colony feeding experiments. Through this multi-pronged investigation, we demonstrated how specialized symbiotic relationships underpin the nutritional ecology of these herbivorous ants.

## Results

### *T. nigra-*group ant workers have evolved an unusual organ — bacterial pouch

Previous studies have revealed the presence of a pouch-like structure in the gut of adult workers from three *T. nigra*-group ant species^32^. Our microscopy analyses of the four ant species from the *T. nigra-*group, including two named^32^ (*T. attenuata*, *T. binghami)* and two so far undescribed (*T.* sp. 1 and *T.* sp. 2 - Fig. 1a and Supplementary Fig. 1), confirmed that this specialized gut structure is consistently present within the group. In adult workers, this pouch-like structure is located at the junction between the midgut and hindgut and is connected to several Malpighian tubules (Fig. 1c, i-l). The inner lining of the pouch lumen consists of microvillar cells containing numerous mitochondria, and a dense population of rod-shaped bacteria enclosed by a double membrane is housed inside the pouch (Fig. 1f and g). The whole-mount fluorescence in situ hybridization (FISH) reveals bacteria are concentrated within the pouch-like structure in all four ant species (Fig. 1i--l), confirming its role as a symbiotic organ, referred to as the bacterial pouch in the *T. nigra-*group ants.

When comparing across life stages of *T. nigra-*group ants, scanning electron microscopy observations revealed that larvae lack the specialized bacterial pouch (Fig. 1e), which only develops during the pupal stage (Fig. 1d). The symbiotic organ in the pupal stage is underdeveloped compared to that of adults: smaller in size and not yet fully swollen. Furthermore, the results from the transmission electron microscopy showed a notable absence of bacteria in the symbiotic organ during the pupal stage (Fig. 1h). This indicates that the establishment of symbiotic relationships occurs after worker eclosion.

### Microbiota of *T. nigra-*group adult ants are consistently dominated by pouch-associated Hyphomicrobiales bacteria

16S rRNA amplicon sequencing for dissected gasters of workers from 4-6 colonies from each of the four *T. nigra-*group species revealed striking consistency in bacterial communities (Fig. 2a). In all individuals from across the surveyed colonies of the four species, the microbiota was predominantly composed of a single OTU assigned to the order Hyphomicrobiales (formerly Rhizobiales), representing between 61.1% and 100% of all reads (median: 99.39%). Each ant species harbored a single distinct Hyphomicrobiales zOTU. In addition, we identified an OTU representing unclassified members of the Alphaproteobacteria class in three species. One of their zOTUs was present in all individuals of *T.* sp. 1 and *T.* sp. 2, and the other was found in all individuals of *T. binghami.* Despite consistent presence, their relative abundance was relatively low (e.g., 0.1–1.1% in *T.* sp. 1, 0.06–3.4% in *T.* sp. 2, 0.3–11.5% in *T. binghami*). Furthermore, we found that all *T. binghami* individuals hosted an additional zOTU from the order Xanthomonadales, with relative abundances ranging from 1.4% to 27.7%. All other bacterial zOTUs combined comprised an average of 0.13% of reads per library (ranging from 0 to 2.55%).

**Figure 2.**
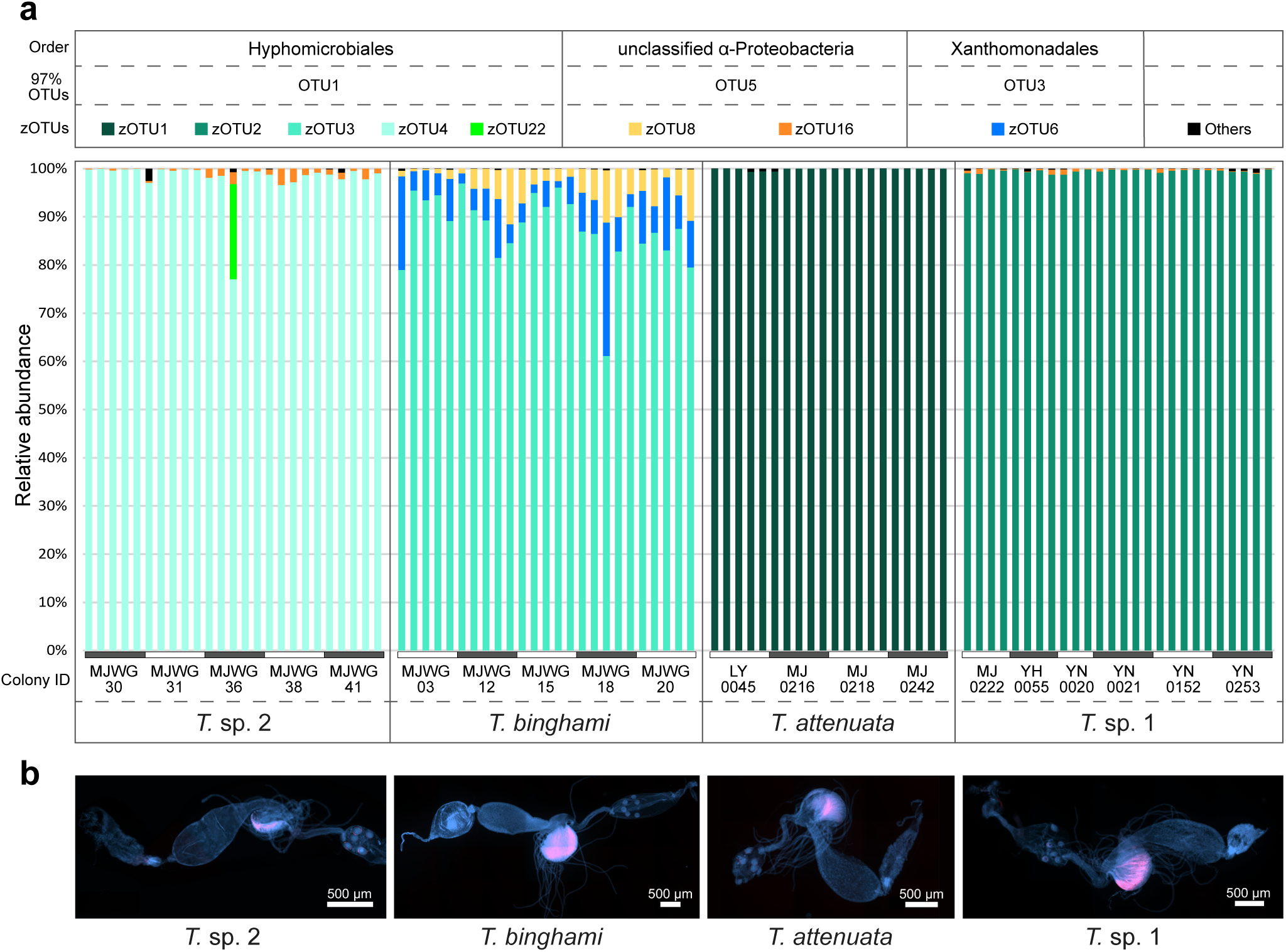
The microbial diversity and the localization of the dominant Hyphomicrobiales bacteria within the gut of workers from four *T. nigra*-group species. **(a)** The relative abundances of bacterial zOTUs across individual workers from replicate colonies of four ant species. zOTUs with a relative abundance of less than 1% in any sample and the prevalence of less than 80% occurrence in any ant species are summed up as “others”. **(b)** FISH microscopy illustrates the localization of Hyphomicrobiales bacteria within the worker ants’ guts. Blue, DAPI; Red, Hyphomicrobiales-specific probe.

We used fluorescence in situ hybridization, with probes specific to each Hyphomicrobiales zOTU to localize Hyphomicrobiales in worker guts of four ant species. Notably, the signal exhibited complete spatial overlap with those of universal bacterial probes, demonstrating that Hyphomicrobiales dominantly colonize the bacterial pouch (Fig. 2b). Diagnostic PCRs targeting Hyphomicrobiales in different life stages of *T. attenuata* and *T.* sp. 1 indicated that Hyphomicrobiales was exclusively present in worker ants, with no detectable presence in larvae or pupae (Supplementary Fig. 2).

### Comparative genomics reveals conserved symbiont lineages in the *T. nigra-*group

After sequencing metagenomes for pooled worker guts from the four *T. nigra-* group species, we performed metagenomic binning to generate draft symbiont genomes, all with more than 89% completeness and less than 1% contamination from the gut microbiomes of four ant species according to CheckM (Supplementary Table 1). The draft genome classification through comparison with the Genome Taxonomy Database aligned with the 16S rRNA amplicon-based findings (Fig. 3a). Specifically, all *Tetraponera* species hosted a dominant and unique Hyphomicrobiales strain. Additionally, *T. binghami* and *T.* sp. 1 were found to carry an unclassified Alphaproteobacteria strain, while *T. binghami* uniquely possessed a distinct Xanthomonadales strain.

**Figure 3.**
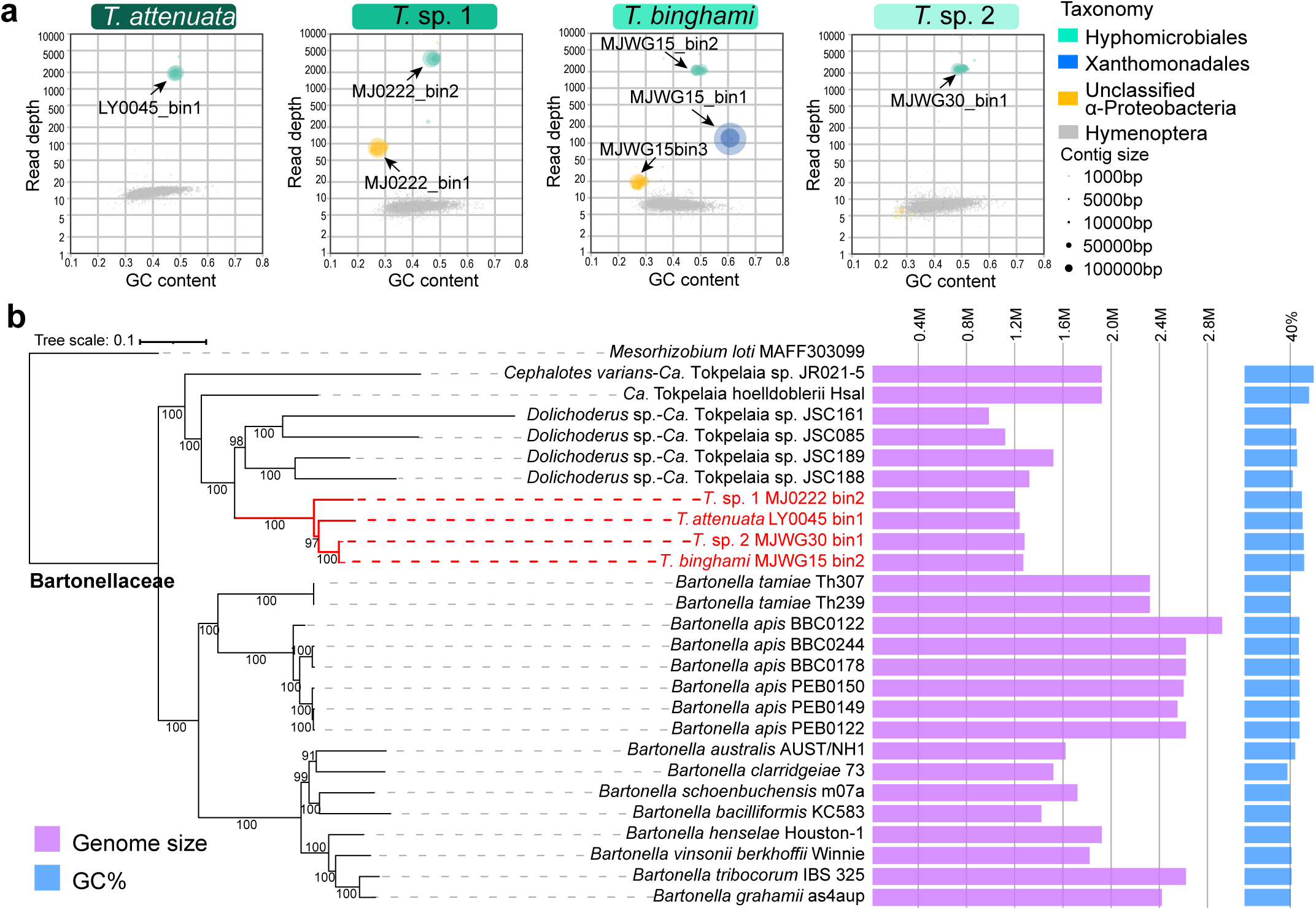
Genomic and phylogenetic characterization of *Tetraponera*-associated symbionts. **(a)** Taxon-annotated GC-coverage plots for four *Tetraponera* metagenomes. Each point presents a scaffold positioned by %GC content (x-axis) and sequencing depth (y-axis, log scale), with color indicating taxonomic assignment (unclassified scaffolds omitted) and size reflecting scaffold length. **(b)** Phylogenomic analysis of dominant Hyphomicrobiales symbionts. Maximum likelihood analysis was performed based on 385 homologous single-copy genes. Red branches represent the Hyphomicrobiales genomes obtained from the metagenome of four *T. nigra*-group ant species. Purple and blue bars represent genome size and GC content, respectively.

The draft genomes of Hyphomicrobiales from the four ant species ranged in size between 1.18 and 1.26 Mb, with an average GC content between 47.4% and 49.5%. The maximum likelihood phylogeny based on a concatenated alignment of 385 amino acid sequences of single-copy orthologous genes confirmed that the four Hyphomicrobiales symbionts grouped together and formed a distinct, well-supported monophyletic clade, along with other ant-associated Hyphomicrobiales symbionts with sequenced genomes (Fig. 3b). This ant-associated bacterial clade forms a sister group to the *Bartonella* clade, which includes *B. apis* of honeybees and multiple mammalian pathogens. This result is also supported by the phylogenetic tree constructed based on 16S rRNA sequences (Supplementary Fig. 3). Intriguingly, the phylogenomic relationships among bacterial strains did not fully agree with those of their ant hosts: strains from *T. attenuata* and *T.* sp. 1 were not identified as sister clades, unlike their hosts. On the other hand, genome organization comparisons revealed that despite multiple rearrangements, there are large areas of synteny among symbiont genomes of *T. nigra-*group ants – much more so than relative to *Tokpelaia* genomes from other ants (Supplementary Figs 4-5) - further supporting the monophyly of this clade. The 16S rRNA gene identities and the average amino acid identities in all pairwise comparisons between all ant-associated Hyphomicrobiales symbionts were 94.61%-97.91% and 62.33%-97.23% (Supplementary Table 2), respectively, suggesting that they should be considered as members of the same genus *Tokpelaia*. Considering this, and the phylogenomic evidence for the single origin of this symbiosis, we propose to consider the *T. nigra-*group symbionts as strains of a single species, *Candidatus* Tokpelaia tetraponerae.

Our 16S rRNA gene phylogeny (Supplementary Fig. 3) indicated that the *T. nigra–* group associated unclassified Alphaproteobacteria and Xanthomonadales symbionts were grouped into well-supported *Tetraponera*-specific clades. Metagenomic binning helped uncover the genomic characteristics of these bacteria. The unclassified Alphaproteobacteria strains had genomes of approximately 1.13 Mb with a GC content of 27.4% (Supplementary Table 1). Phylogenetic analysis placed them within the order Holosporales (Supplementary Fig. 6), a group known to include obligate intracellular or intranuclear symbionts of protists^39,40^. In addition, the Xanthomonadales bacterium, found exclusively in *T. binghami*, had a genome size of 2.36 Mb and a GC content of 60.99% (Supplementary Table 1). Phylogenetic analysis revealed its close relationship to *Frateuria aurantia* isolated from a lily plant, *Lilium auratum.* However, taxonomic inconsistencies within the *Frateuria* genus preclude precise genus-level assignment, and thus, we provisionally classify it within the Rhodanobacteraceae family. While this last symbiont has the largest genome among the symbionts of *T. nigra-*group ants, it is still reduced by one-third compared to the 3.6 Mb genomes of closely related bacteria (Supplementary Fig. 6).

### Nitrogen recycling and provisioning facilitated by the *T. nigra-*group ant symbionts

Given that the diets of these ants are deficient in nitrogen, yet their symbionts reside in a bacterial pouch enriched with nitrogenous waste such as uric acid delivered through the Malpighian tubules, we proposed that these symbionts may participate in nitrogen recycling. Analysis of *Tokpelaia* genomes supports this hypothesis, as they retained genes encoding enzymes involved in nitrogen cycling from nitrogenous waste and biosynthetic pathways for the production of most essential amino acids that are needed by the ant hosts (Fig. 4).

**Figure 4.**
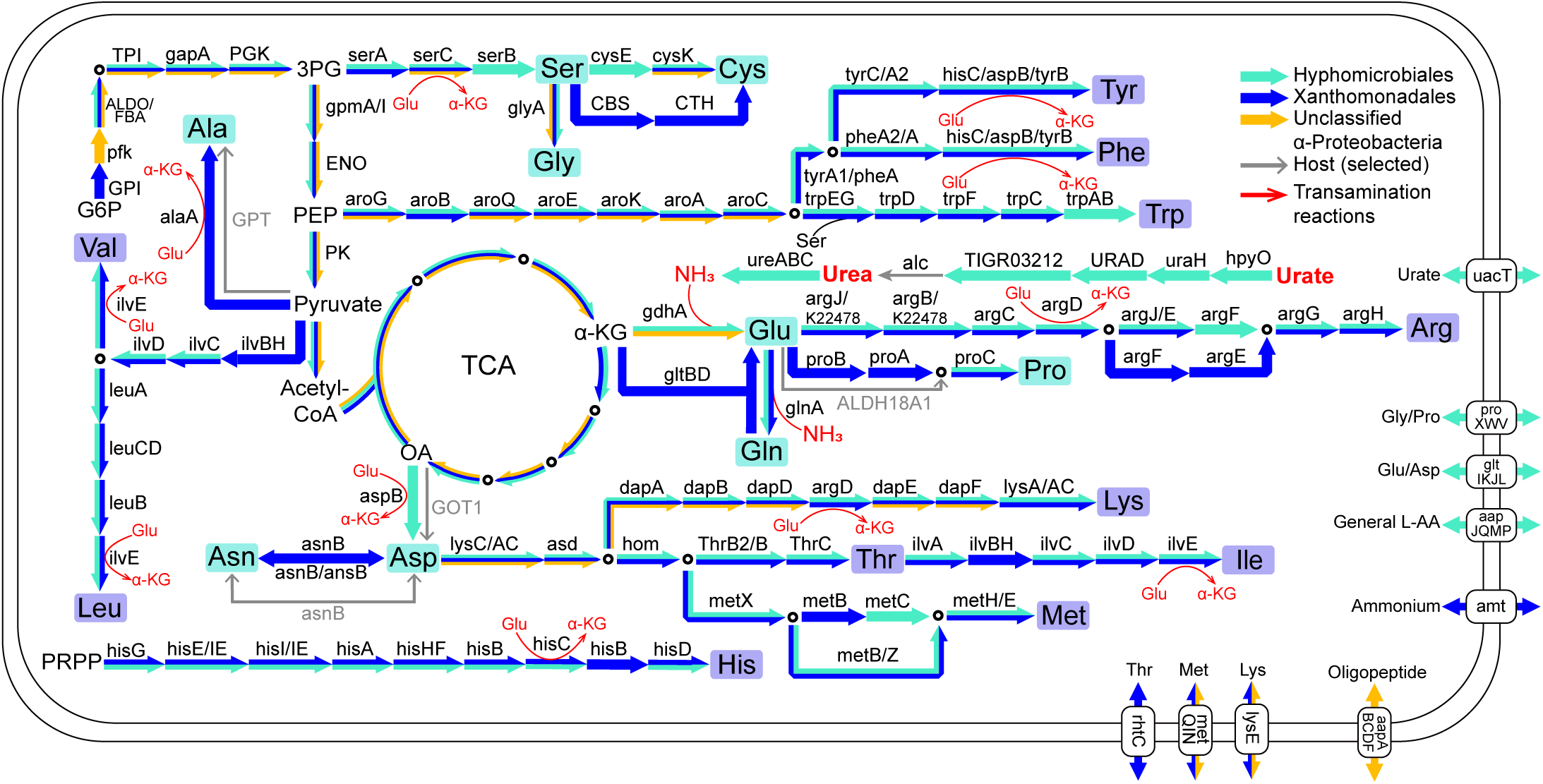
Metabolic reconstruction of nitrogen waste recycling and amino acid biosynthesis pathways in *Tetraponera*-associated symbiont genomes. Pathways were predicted from draft genomes of symbiont strains obtained from four *Tetraponera* metagenomes. Arrow colors indicate contributions from distinct bacterial symbionts or the ant host. Key metabolic intermediates are abbreviated: G6P, Glucose 6-phosphate; 3PG, 3-phosphoglycerate; PEP, phosphoenolpyruvate; PRPP, phosphoribosyl pyrophosphate; OA, oxaloacetate; α-KG, alpha-ketoglutarate. Red arrows denote transamination reactions, where an amino group from glutamate are donated to the given amino acid precursors.

All the *Tokpelaia* genomes harbored the *uacT* gene, which encoded a transporter for uric acid—the primary metabolic waste transported by the host’s Malpighian tubules into the bacterial pouch. Further, all *Tokpelaia* strains contain gene clusters encoding enzymes involved in all the uric acid degradation pathway steps except for the final reaction (Fig. 4). Interestingly, the missing gene, *alc, which* encodes allantoicase that converts allantoate to urea, seems ancestrally lost in *Tokpelaia*, but it is present on Hymenoptera-assigned scaffolds in metagenomes of the *T. nigra-*group ants except *T.* sp. 1, implicating that this reaction is likely carried out by the ant hosts (Supplementary Fig. 7). Following uric acid degradation, the resulting urea could be further processed into ammonia through the urease, encoded by three core genes, *ureC*, *ureB,* and *ureA* (Supplementary Data 2), present in all sampled metagenomes. Further, all *Tokpelaia* strains possess the genes *gdhA* and *glnA*, encoding enzymes glutamate dehydrogenase (GDH) and glutamine synthetase (GS), respectively (Fig. 4). These enzymes assimilate ammonia – like that from urease-driven urea degradation – producing glutamate and glutamine, which serve as the nitrogen donors for the biosynthesis of amino acids.

The *Tokpelaia* genomes retain complete pathways for the biosynthesis of the essential amino acids phenylalanine, tryptophan, tyrosine, arginine, lysine, threonine, and methionine (Fig. 4, Supplementary Fig. 8). The isoleucine, leucine, and valine pathways are nearly complete, although *ilvH* gene encoding the small subunit of acetolactate synthase is missing. It is possible that in *Tokpelaia* these reactions are catalyzed solely by the large subunit (IlvB) of acetolactate synthase, as demonstrated in some Enterobacteria bacteria^41^.

Despite their specialized role in nitrogen metabolism, the four *Tokpelaia* strains exhibit striking metabolic constraints. While retaining genes for core cellular processes, including translation, replication, and transcription, their metabolic gene repertoire is notably limited (Supplementary Figs. 9-11). The symbionts’ genomes lacked key genes involved in glycolysis, the biosynthesis of most B vitamins, certain non-essential amino acids, nucleotides, and KDO_2_-lipid A. However, the metabolic pathways of these symbionts remain intact for the TCA cycle, oxidative phosphorylation, the biosynthesis of peptidoglycan, and fatty acids.

Analysis of the Xanthomonadales genome uniquely found in *T. binghami* revealed its metabolic redundancy, as it harbored genes encoding enzymes involved in ammonia assimilation and the biosynthesis of most amino acids (Fig. 4). Particularly, while genes *ilvH* and *hisB* were missing from across all examined *Tokpelaia* genomes, both were present in the Xanthomonadales genomes. In contrast, Holosporales strains lack biosynthesis pathways for almost all amino acids (Fig. 4, Supplementary Fig. 8), but encode transporters for oligopeptides as well as several amino acids. In addition, they also harbor glucose PTS transporters encoded by ptsG, enabling glucose to serve as the substrate fueling their TCA cycle (Supplementary Data 2). Like other Holosporales bacteria reported previously^42,43^, these bacteria are unlikely to be involved in ant nutrition, despite their strikingly consistent presence.

### Microbiota disruption negatively affects the colony-wide fitness

Given the metagenomic evidence for nutritional provisioning by pouch-associated symbionts, we assessed symbiont impact on host survivorship and development. We first generated symbiont-free workers by removing pupae from the original colonies to sterile dishes. These isolated pupae kept under sterile conditions emerged as adult workers with no bacteria in the bacterial pouch, confirmed by universal bacterial PCRs, diagnostic *Tokpelaia* 16S rRNA gene-targeting PCRs, and whole-mount FISH staining (Supplementary Fig. 12).

We then compared fitness parameters between the two groups of larvae with similar sizes: one reared by normal workers (Sym-workers) and the other reared by symbiont-free, aposymbiotic workers (Apo-workers), in two ant species, *T. attenuata* and *T.* sp. 1. Through detailed daily observation of larval development status over a 30-day period, we found that larvae tended by symbiont-free workers in both species exhibited significantly lower survivorship (Fig. 5a; Wald statistic = 192.6, p<2e-16 for *T.* sp. 1; Wald statistic = 180.6, p <2e-16 for *T. attenuata*) and slower development, as indicated by reduced body length (Fig. 5b; χ² =897.982, p <2.2e-16 for *T.* sp. 1; χ² =721.6, p <2.2e-16 for *T. attenuata*; GLMM) compared to larvae reared by normal workers.

**Figure 5.**
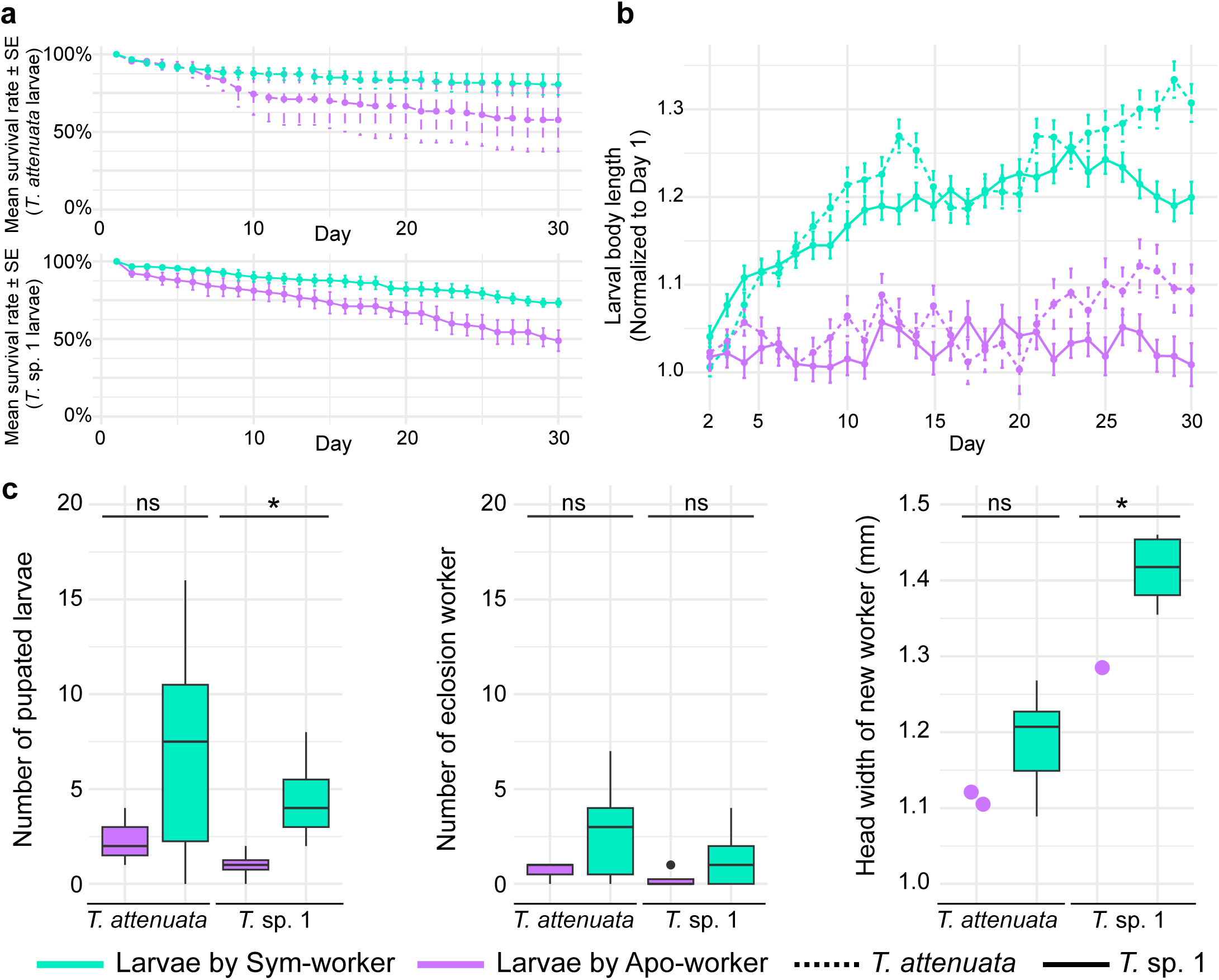
Comparison of fitness parameters between two groups of larvae with similar sizes, one reared by normal workers (Sym-workers) and the other reared by symbiont-free workers (Apo-workers), in two ant species, *T. attenuata* and *T.* sp. 1. **(a)** The survival rate of larvae cared for by Apo-workers or Sym-workers over a 30-day experimental period. **(b)** Changes in the relative body length of larvae reared by Sym-workers and Apo-workers. Body length data from day 2 to day 30 were normalized to the average body length of larvae on day 1 for each treatment group. **(c)** Comparison of fitness parameters at day 90, including pupation rate, eclosion rate, and head width of newly emerged workers as an indicator of body size. The light green color represents the control group of worker ants with symbiotic bacteria, while the purple indicates the experimental group of worker ants without symbiotic bacteria. In the line graphs, the dashed lines represent *T. attenuata*, and the solid lines represent *T.* sp. 1.

In a separate 90-day experiment, we monitored the continuous development of the larvae until they emerged as adults. After 90 days, larvae raised by symbiont-free workers exhibited lower pupation rates (Fig. 5c, Supplementary Table 3; *T.* sp. 1, p = 0.0096; *T. attenuata*, p = 0.2326; T-test), reduced adult emergence rates from pupae to adults (Fig. 5c, Supplementary Table 3; *T.* sp. 1, p = 0.1999, Wilcoxon-Mann-Whitney test; *T. attenuata*, p = 0.2271, T-test), as well as smaller head widths at the time of emergence (Fig. 5c, Supplementary Table 3; *T.* sp. 1, p = 0.1818, Wilcoxon-Mann-Whitney test; *T. attenuata*, p = 0.0453, T-test), compared to those reared by normal workers. We also noted that among the adult ants tending the larvae, symbiont-free workers exhibited lower survivorship compared to their symbiont-bearing counterparts (Supplementary Fig. 13; Wald statistic = 94.22, p<2e-16 for *T.* sp. 1; Wald statistic = 3.44, p = 0.06 for *T. attenuata*).

### Symbionts recycle nitrogen from urea and provide it to ants

Building on metagenomic evidence for symbiont-encoded ability to degrade nitrogenous wastes and synthesize amino acids, we conducted feeding experiments with ^15^N-labelled urea (Supplementary Fig. 14) on two species from the *T. nigra-*group. Within each ant colony, individuals were assigned to one of three experimental groups: (1) symbiotic workers fed ^15^N-labelled urea; (2) symbiont-free workers fed ^15^N-labelled urea; and (3) symbiont workers fed ^14^N-labelled urea as a control.

In the first experiment, we focused exclusively on *T. attenuata* workers from three colonies. We examined whether recycled nitrogen from ^15^N-labelled urea could be incorporated into the free amino acids present in the hemolymph of adult ants. Among symbiont workers fed ^15^N-urea, 14 of the 19 measurable amino acids were enriched for the heavy ^15^N isotope compared to the control group fed diets containing standard, ^14^N-labelled urea (significant p-value range: 1.04E−05 to 0.076, Fig. 6a, Supplementary Table 4). They included seven essential amino acids (arginine, histidine, isoleucine, phenylalanine, tryptophan, tyrosine, and valine), as well as seven non-essential amino acids (alanine, asparagine, glutamate, glutamine, glycine, proline, and serine). Crucially, this heavy isotopic enrichment was dependent on symbiont presence. Symbiont-free *T. attenuata* workers fed ^15^N-labelled urea showed no such enrichment in the same amino acids relative to the same control. These results highlighted the pivotal role of bacterial pouch symbionts in recycling nitrogen, particularly in providing amino acids essential to the hosts.

**Figure 6.**
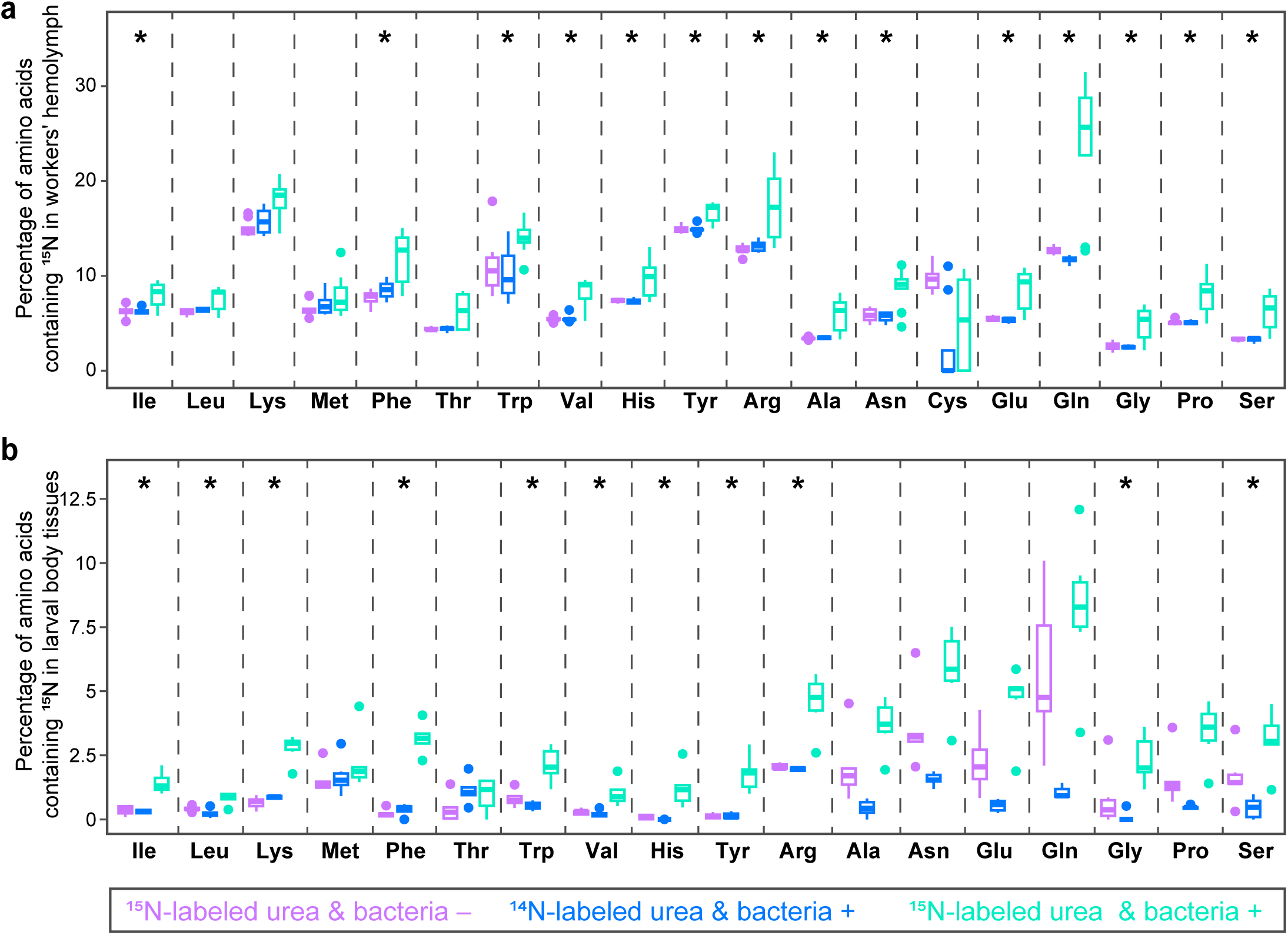
Depriving symbiotic bacteria reduces the proportion of 15N-labeled amino acids in the worker ants and larvae that have consumed 15N-labeled urea. **(a)** The proportion of 15N-labeled amino acids in the hemolymph of *T. attenuata* workers was measured under three conditions: symbiotic workers fed 15N-labeled urea, symbiont-free workers fed 15N-labeled urea, and symbiotic workers fed 14N-labeled urea as a control. Detailed information on sample sizes for each treatment can be found in Supplementary Data 10. The statistical results are described in the Supplementary Methods and Supplementary Table 9. **(b)** The proportions of 15N-labeled amino acids in the body tissues of larvae cared for by workers of *T. attenuata* and *T.* sp. 1 under the same three conditions. Due to limited sample size per species (n=3 under one condition) and non-significant interspecific effect on the proportion of 15N amino acids (Supplementary Table 11), data from two species were pooled across group for a combined analysis using one-way ANOVA (for normally distributed data) or Kruskal Wallis test (for non-normal data).

In a follow-up experiment, we investigated how bacterial pouch symbionts coordinate colony-wide nitrogen provisioning. We implemented two tracing approaches by including larvae tended by adult workers in *T. attenuata* and *T.* sp. 1 system. To delineate the metabolic contribution of symbionts to larval nutrition, we first tracked ¹⁵N incorporation into amino acid pools in body tissues of larvae tended by workers in the ¹⁵N-labeled urea feeding experiments. We found that larvae reared for 30 days by symbiotic workers fed ^15^N-urea exhibited ^15^N enrichment in 11 of the 18 measurable amino acids, relative to larvae fed by workers provided ^14^N-urea (p-value ranges: 0.001 to 0.0861, Fig. 6b, Supplementary Table 5). These 11 enriched amino acids included nine essential amino acids (arginine, histidine, isoleucine, leucine, lysine, phenylalanine, tryptophan, tyrosine, and valine) and two non-essential amino acids (glycine and serine). In contrast, larvae tended by symbiont-free workers (^15^N-urea fed) showed no such enrichment. This highlights the worker symbionts’ roles in bridging nutritional needs critical for larval growth and development.

During the same experiment, we also quantified bulk δ¹⁵N in both larval and worker bodies to assess how recycled nitrogen is partitioned between castes. We observed that δ^15^N values in the body tissues of larvae attended by symbiotic workers for 30 days were significantly higher compared to those reared by aposymbiotic workers (*T. attenuata*, p=0.0373; *T.* sp. 1, p=0.0399; post hoc Dunn’s test; Supplementary Fig. 15, Supplementary Table 6). Similarly, δ^15^N values in the body tissues of symbiotic workers were moderately higher than those of aposymbiotic workers (*T. attenuata*, p=0.1062; *T.* sp. 1, p=0.1052; post hoc Dunn’s test; Supplementary Fig. 15, Supplementary Table 6). Strikingly, comparative analysis revealed that larvae exhibited 2.2-fold and 2.3-fold higher 15N enrichment compared to symbiotic workers tending them in *T. attenuata* and *T.* sp. 1, respectively (*T. attenuata*, p=0.0022; *T.* sp. 1, p=0.0005; t-test). These findings highlight how microbial symbioses help ants address colonies’ nutritional needs with emphasis on the critical larval stage, thus maximizing fitness under N-limiting conditions.

## Discussion

Our study has demonstrated the unique location of symbiotic bacteria within a gut pouch (Fig. 1h-1k), a special structure consistently presents in adult ants across the *T. nigra-*group. We also demonstrated for the first time these ants’ ancestral symbioses with *Tokpelaia* symbionts and consistent associations with some other bacteria (Figs. 2 and 3). Through the characterization of the symbionts’ genomes, we have elucidated the mechanisms underlying this nutritional symbiosis (Fig. 4). Additionally, we have demonstrated experimentally the effects of symbiosis disruption on colony development, highlighting the impact on larval growth (Fig. 5). Finally, we have experimentally established links between symbiont genomic characteristics and ant nutrition, explaining the observed fitness effects (Fig. 6). Considering the diversity, global distribution, and ecological importance of social insects, it is surprising how few clades and species have comparably detailed data on the nature and significance of microbial associations. Through this study, the *T. nigra-*group ants are added to the short list of social insect clades whose symbioses are considered well-understood.

The *T. nigra-*group is distinguished by a unique bacterial pouch—a structure not previously reported in other hymenopteran clades. Other hymenopterans exhibit diverse but clearly distinct anatomical niches for hosting symbiotic bacteria. For instance, *Blochmannia* resides in bacteriocytes localized among midgut cells and in female ovaries^44,45^. Species such as honeybees^46,47^, bumblebees^48^, or *Cephalotes* ants^15,49^ harbor multi-partite bacterial communities within the gut chambers. Moreover, fungus-farming ants maintain bacterial communities in specialized exoskeletal crypts essential for the health of their fungal gardens^50^.Beyond hymenopteran insects, the most similar structures were described from stinkbugs and leaf beetles^7,33,51,52^, which host extracellular bacteria in a symbiotic organ consisting of pouch-like gut invagination. These symbiotic organs ensure a stable supply of resources for the symbiotic bacteria, promoting their growth while limiting competition with other microbial species^53^. Meanwhile, these structures restrict the entry of symbionts into other host tissues, minimizing challenges such as the immune response.

While a pouch-like structure in the gut of adult *T. nigra*-group workers had been described previously^32,54^, we demonstrate for the first time that this specialized anatomical structure exhibits life stage specificity. Absent in larvae, the pouch first develops during the pupal stage, but it remains uncolonized until the ants eclose as adults. This indicated that this symbiotic organ develops independently of symbiont colonization. A similar pattern of organ development, proceeding without symbiont triggering, has been observed in tortoise beetles, reflecting an ancient evolutionary trait in which the formation of such organs proceeds autonomously^55^. Moreover, bacterial colonization of their pouch depended on contact with colony members, suggesting that symbiotic bacteria are transmitted to newly emerged adults through trophallactic interactions. Such trophallaxis behaviors may play a key role in maintaining partner fidelity, not only in these groups of ants but also across several other social insects. For instance, symbionts are transmitted between termite siblings through oral-anal trophallaxis^56^, and a similar transmission mode has been observed in *Cephalotes* and *Procryptocerus* ants^57^. In honeybees, gut symbiotic bacteria are transmitted through a combination of trophallaxis and contact with nest materials within the hive^58^.

High-throughput sequencing revealed an ant-specific, *Tokpelaia*-dominated bacterial community within the pouch, suggesting a specialized and long-standing association between the *T. nigra-*group ants and their symbiotic bacteria. Our conclusions are further strengthened by the phylogenetic similarity between the symbionts characterized here and 16S rRNA data previously generated from other *T. nigra-*group species^54,59^. Our findings suggest that the *T. nigra-*group ants belong to a growing number of insect clades whose extracellular microbiota are dominated by one or a few specialized symbionts—a pattern similarly observed in other insect groups with specialized symbiotic organs, such as stinkbugs^7^ and leaf beetles^33,51,60^. One possible explanation is that these symbiotic organs function to filter microbial strains, allowing only specific symbionts to access the host and maintain a beneficial partnership. This contrasts with the complex multi-partite gut microbiota observed in termites^61,62^, *Cephalotes* ants^15^, and corbiculate bees^47^, or with occasional and non-specialized microbial association seen in other ants and other insects^63^.

Symbiotic bacteria play a critical role in the colony-wide fitness of *T. nigra*-group ants. This is reflected by higher survival of ant workers retaining symbionts, improved survival and growth of the larvae, and substantial ^15^N enrichment of amino acid pool and body tissues in both adults and larvae, with larvae exhibiting a higher rate of nitrogen incorporation but only when microbiota were undisturbed. This highlights the contribution of symbiont-mediated nitrogen metabolism to the fitness of the ant colony. Moreover, genomic analysis and heavy nitrogen urea labeling experiments elucidated the underlying mechanisms: symbionts in the bacterial pouch recycle nitrogenous waste, such as urate or urea, and convert symbiont-recycled nitrogen in the form of essential and non-essential amino acids utilized by ant hosts. Taken together, this work provides robust evidence supporting the 23-year-old^54^ of the conserved N-recycling roles of symbionts in the bacterial pouch of *T. nigra-*group ants, revealing that this nutritional symbiosis plays an important role for ant hosts in the exploitation of an N-limited dietary niche. Similar functions are observed in other herbivorous ants, such as the endosymbiont *Blochmannia* in Camponotini and the multi-partite gut bacterial communities in *Cephalotes* ants. Despite differences in the nature of microbial symbioses, these ants with nitrogen-limited diets exhibit a functional convergence in the adaptive role of symbiosis for nitrogen acquisition. This reflects the generality of such functional convergence across herbivorous ant species inhabiting N-limited environments.

The specialized structure connecting the bacterial pouch to the Malpighian tubules reinforces the proposed biological role of *Tokpelaia*. The nitrogen recycling activity by *Tokpelaia* is likely mediated through the delivery of host-derived waste uric acid via the Malpighian tubules into the bacterial pouch where *Tokpelaia* resides. Additionally, this structure’s connection to the gut facilitates the export of amino acids synthesized from assimilated recycled nitrogen by *Tokpelaia* cells, which are then transported to the host gut. The high abundance of nitrogen-recycling symbionts at the site of insect nitrogen waste delivery via the Malpighian tubules is also observed in the *Cephalotes* system. Positioned at the site of host nitrogen waste delivery^15,49^, *Ischyrobacter davidsoniae* and other bacteria play a central role in a collaborative nitrogen-recycling symbiosis^23^. These findings highlight the critical role of specialized symbiotic structures in facilitating nitrogen recycling in insects, where the spatial arrangement of symbionts near nitrogen waste delivery sites may enable efficient nutrient cycling and contributes to the overall ecological success of the host.

During the metamorphosis of holometabolous insects, significant morphological and physiological changes provide opportunities for symbiotic mutualisms to become dissociated. In social insects, such cross-stage symbiont benefits are more pronounced due to the close physical association between adults and larvae and the capacity for trophallactic feeding of symbiont-derived metabolites produced by adults. In the *T. nigra-*group ants, specialized *Tokpelaia* symbionts, present exclusively in adults, enhance colony-wide nitrogen economies. Their contributions to larval nitrogen budgets are likely to promote larval growth and development. This adult-specific symbiosis is likely sustained through the trophallactic transfer of extracellular symbionts during the adult stage.

Taken together, our findings indicate that the bacterial pouch – an unusual symbiotic organ that might be unique in Hymenoptera - enables an efficient nitrogen economy within the holobiont of *T. nigra-*group ant species. The compartmentalized structure suggests a potential mechanism for ant hosts to deliver precursors of the uricolytic pathway directly to the bacterial pouch, though further validation is needed. The symbionts, which reside within this structure, recycle nitrogenous waste either through host metabolism or the host diet and produce metabolites that are beneficial to the ants, functioning as “nutrient factories”. This system defines, for the first time, the metabolic capabilities of *T. nigra-*group ants’ symbionts, representing a striking example of convergent evolution among herbivorous ants, yet with a unique morphological innovation. Crucially, these symbiont-derived metabolites are then passed on to colony members, particularly larvae that lack these symbionts, via trophallaxis, ensuring that the entire colony benefits from the nutritional symbiosis. This out-of-body nutritional symbiosis, where symbiont benefits transcend individual hosts to support colony-wide fitness, represents the first demonstration of cross-caste nutritional integration in social insects. Our findings redefine how symbiotic partnerships can be integrated into superorganismal biology, offering new insights into the evolutionary mechanisms underlying ecological success in nutrient-poor environments.

## Methods

### Ant collection and specimen identification

We collected 53 *Tetraponera* colonies from southern China (Yunnan, Guangdong, and Jiangxi) between 2019 and 2024 (Supplementary Data 1). For molecular analysis, workers were preserved in anhydrous ethanol at −20°C before DNA extraction, while live colonies were maintained for dietary manipulation experiments or microscopic observation. Species identification combined morphological classification and mitochondrial COI barcoding. COI sequences (>600 bp) were aligned with reference sequences from GenBank and BOLD databases, followed by phylogenetic analysis (maximum likelihood with 1000 bootstrap replicates) and species delimitation via the Automatic Barcode Gap Discovery (ABGD) method^64^ (Supplementary Fig. 1).

### Trophic levels of *Tetraponera* ants determined by nitrogen isotope ratios

We collected local plants, herbivores, and predatory arthropods alongside *Tetraponera* ants for stable isotope analysis. All samples were transported to the laboratory, where arthropod abdomens were removed to eliminate potential gut content contamination. Samples were dried, ground, and analyzed for δ¹⁵N values (Supplementary Data 3) using a Delta A Advantage isotope ratio mass spectrometer coupled with an EA-HT elemental analyzer (Thermo Fisher Scientific Inc., Bremen, Germany). Trophic positions were calculated by: (1) establishing site-specific plant δ¹⁵N baselines, (2) computing Δδ¹⁵N values (δ¹⁵Nconsumer - δ¹⁵Nplant) for each ant and arthropod specimen, and (3) applying regional corrections by normalizing to median plant values across all sites.

### Microscopic analysis of bacterial pouch structure and symbiont localization in *T. nigra-g*roup Ants

To characterize the bacterial pouch organization across developmental stages in *T. nigra*-group ants, we employed complementary microscopy approaches. First, scanning electron microscopy (SEM) was used to examine the gut structure and bacterial pouch morphology in workers, pupae, and larvae of *T. attenuata*. Fluorescence in situ hybridization (FISH) with EUB338 universal bacterial probe and Rhizobiales-specific probes were then used to localize symbiotic bacteria within worker guts across four *T. nigra*-group ant species. Finally, transmission electron microscopy (TEM) provided ultrastructural details of the bacterial pouch and associated bacteria in *T. attenuata*. Detailed protocols are provided in the Supplementary Methods.

### Bacterial 16S rRNA amplicon sequencing

We characterized the gut microbial communities of *T. nigra*-group ants through the V4 region of the 16S rRNA amplicon sequencing. For each species, 3-5 workers from 4-6 colonies were sampled. DNA was extracted from the dissected gaster of a single worker using a commercial kit (Qiagen), followed by library preparation and paired-end sequencing on Illumina NovaSeq6000 SP platform, generating 2×250bp reads (Supplementary Data 4). Along with insect DNA samples, blank samples in our DNA extraction batches were included to monitor potential contamination during sample preparation and sequencing. Raw reads were first demultiplexed and merged, followed by quality filtering, chimera removal, and denoising to generate zOTU and 97% OTU tables using a modified version of the “modified_LSD.py” script^65^ from https://github.com/catesval/army_ant_myrmecophiles. Taxonomic classification was performed against RDP databases. We implemented stringent contamination filtering using blank controls and cloned reference sequences to generate the quality-filtered zOTU table (Supplementary Data 5). Complete protocols are available in the Supplementary Methods.

### Metagenomics

For each ant species, we dissected the gut tissues of 10 workers from the same colony using sterile forceps and pooled them into a single sample. The extracted DNA was then sent to Beijing Novogene Bioinformatics Technology Co., Ltd for library preparation, followed by metagenomic sequencing on the Illumina NovaSeq 6000 platform. Quality trimming of raw reads was done with Trimmomatic-0.39^66^, followed by read quality checks using FastQC v0.11.9(http://www.bioinformatics.babraham.ac.uk/projects/fastqc). The trimmed reads were assembled with IDBA-UD 1.1.3^67^ using k values of 20, 40, 60, 80, and 100, and assembly statistics were calculated with QUAST-5.0.2^68^ (Supplementary Table 7). Scaffolds were uploaded to the Integrated Microbial Genomes with Microbiome Samples Expert Review (IMG/M-ER) website for taxonomic classification and gene annotation^69,70^. We focused on N metabolism by utilizing KEGG as references to manually construct the pathways involved in the degradation of nitrogenous wastes and the biosynthesis of amino acids (Supplementary Data 2). The bacterial community composition of all metagenomes in this study was visualized by the Taxon-Annotated GC Coverage plots, which employed custom scripts based on a previously published method^71^.

We obtained the draft genomes of individual symbiont strains using the metaWRAP pipeline^72^. Draft genomes were refined with the bin_refinement module using ≥80% completeness and ≤10% contamination. Scaffolds with unexpected taxonomic classifications or inconsistent coverage were manually removed following comparison with the NCBI database using BLASTn and BLASTx (Supplementary Data 6). Bin completeness and contamination were assessed with CheckM^73^ and the taxonomic assignment was performed using the Genome Taxonomy Database with GTDB-Tk v2.1.1^74^ (Supplementary Table 1).

### Phylogenetic analysis of symbiotic bacteria

We reconstructed the evolutionary relationships of dominant bacterial symbionts in *T. nigra*-group ants through two phylogenetic approaches. For 16S rRNA-based analysis, we selected high-abundance zOTUs (relative abundance >0.01 in ≥80% of conspecific libraries), as well as sequences obtained through cloning and metagenomic sequencing. Reference sequences were selected through BLASTn searches, retaining ant-associated lineages and three non-ant associates from top BLAST hits. We further performed genome-based phylogenetic analysis of symbiotic bacteria in *T. nigra*-group ants. We identified single-copy orthologous genes from symbiont draft genomes and other selected genomes listed in Supplementary Data 7, generated concatenated amino acid sequence alignments, and selected optimal substitution models. Both analyses utilized maximum likelihood approach using RAxML v8.2.12^75^ with 1000 bootstraps. Results were visualized and annotated on iTOL^76^.

### Symbiont manipulation and ^15^N-labeled urea feeding experiments

In order to perform symbiont manipulation experiments, we first obtained large numbers of symbiont-free workers without antibiotics treatment. Pupae with pigmented eyes from *T. attenuata* and *T.* sp. 1 lab-reared colonies were removed from the original cages and placed in sterile dishes. Those pupae kept under sterile conditions will emerge as adult workers with no bacteria in the bacterial pouch, confirmed by PCR amplification using both universal bacterial and *Tokpelaia*-specific 16S rRNA gene primers, as well as whole-mount FISH staining (Supplementary Fig. 12).

Controlled lab experiments were conducted on two ant species, *T. attenuata* and *T.* sp. 1, to quantify the symbionts’ contributions to the colony-wide nitrogen economy through N recycling of dietary N waste (Supplementary Fig. 14). For each species, three or four colonies were selected as biological replicates. Within each colony, individuals were randomly assigned to one of the three treatments: (1) workers harboring symbiotic bacteria in the bacterial pouch (Sym-worker), fed 30% sucrose water containing 1% (w/v) ^15^N-labeled urea; (2) workers without symbiotic bacteria in the bacterial pouch (Apo-worker), fed the same ^15^N-labeled urea solution; (3) workers with symbiotic bacteria in the bacterial pouch, fed 30% sucrose water containing 1% (w/v) ^14^N-labeled urea as a control.

We initially conducted the ¹⁵N-labeled urea feeding experiments exclusively with worker ants for 3 weeks. At the end of the experiment, hemolymph was collected from the surviving workers. Based on the number of remaining workers, we aimed to ensure 2–3 replicates for each treatment group within each colony. Hemolymph from each replicate was pooled to measure the enrichment of ¹⁵N in amino acids within ant hemolymph (Supplementary Data 8) at the Scale Biomedicine Technology Co., LTD (Beijing, China). Chromatographic separation of each hemolymph-water mixture was performed on Ultra-High Performance Liquid Chromatography (UHPLC)–High Resolution Tandem Mass Spectrometry (HRMS/MS).

Next, we extended the ¹⁵N-labeled urea feeding experiments to include larvae tended by adult workers, with experimental durations of 30 days and 90 days. In the 30-day feeding experiment, each treatment group was assigned 10 workers and 30 larvae. At the end of the experiment, δ¹⁵N values in larval and worker body tissues (Supplementary Data 9), excluding the gut to avoid potential food contamination, was measured using a MAT253 stable isotope mass spectrometer at the Beijing Academy of Agriculture and Forestry Sciences (BAAFS). At the same time, 3-5 larvae from each group were sent to Scale Biomedicine Technology Co., LTD (Beijing, China), where the relative abundance of ^15^N-labeled amino acids was analyzed using UHPLC-HRMS/MS (Supplementary Data 10).

In addition, we evaluated colony fitness under different symbiont manipulation treatments. We recorded worker and larval survival rates (Supplementary Data 11) and monitored larval development by measuring larval body length daily (Supplementary Data 12) in the 30-day feeding experiment. We also performed the 90-day experiment, with each group assigned 15 workers and 50 larvae, to record larval pupation and eclosion rates (Supplementary Table 9) and measure the head width of newly emerged workers as an indicator of the worker body size (Supplementary Table 10). The detailed methods and statistical analysis can be found in the Supplementary Methods.

## Acknowledgements

We thank Jacob Russell and Corrie Moreau for helpful comments on an earlier draft of the manuscript. This study was supported by the National Natural Science Foundation of China 32370448 to Y. H., Beijing Advanced Innovation Program for Land Surface Science and the “111” Program of Introducing Talents of Discipline to Universities (B13008).

## Author contributions

M.M.J. and Y.H. conceived the study. M.M.J collected ant colonies. M.M.J and B. R. Z. performed experiments. M.M.J and Y. H. performed data analyses. M.M.J, D.Y.Z, P.L. and Y.H. prepared the manuscript. All authors participated in the manuscript review and editing.

## Data Availability

All data generated in this study are publicly available. The 16S rRNA amplicon sequencing data have been deposited in the GenBank Short Read Archive under BioProject PRJNA1159735, while the 16S rRNA sequences obtained via Sanger sequencing are accessible in GenBank under accession numbers PQ549644-PQ549668. Assembled metagenomes are available in IMG under the following accession numbers: Gp0646313, Gp0688973, Gp0646295 and Gp0754593. Additionally, raw data from the stable isotope mass spectrometer and UHPLC-HRMS/MS are provided in the Supplementary Tables, and details on the PCR primers and FISH probes can be found in the Supplementary Tables and Methods.

