## Supplementary Information for "Symbiotic solutions for colony nutrition: conserved nitrogen recycling within the bacterial pouch of *Tetraponera* ants"

**Description of Supplementary Files**
File name: Supplementary Information
Description: Supplementary Figures, Supplementary Tables, Supplementary Methods and Supplementary Reference.

File name: Supplementary Data 1
Description: Collection information of ant colonies used in this study.

File name: Supplementary Data 2
Description: KEGG Orthologs (KOs) found in seven symbiont draft genomes. Light green highlighted cells correspond to genes illustrated in supplementary figures 8 and 9.

File name: Supplementary Data 3
Description: Information of local plants and arthropods used for trophic level measurements.

File name: Supplementary Data 4
Description: Summary of PCR amplification and library construction for samples submitted for amplicon sequencing

File name: Supplementary Data 5
Description: zOTU table from *T. nigra-group* ant bacterial community samples.

File name: Supplementary Data 6
Description: Summary of scaffolds assigned to 7 bins in four *T. nigra-group* ant metagenomes.

File name: Supplementary Data 7
Description: Information of bacterial genomes used for phylogenetic analysis.

File name: Supplementary Data 8
Description: Information of samples and isotopic peak abundance of amino acids in the hemolymph of workers in the $^{15}N$-labeled urea feeding experiment (M+N abundance (percentage), where N is the isotope atomic number).

File name: Supplementary Data 9
Description: Description: Information of samples and $\delta^{15}N$ value of the body tissues of workers and larvae cared for by Sym-workers and Apo-workers in a 30-day feeding experiment.

File name: Supplementary Data 10
Description: Description: Information of samples and isotopic peak abundance of amino acids from whole larvae reared by Sym-workers and Apo-workers over a 30-day period in a $^{15}N$-urea feeding experiment (M+N abundance (percentage), where N is the isotope atomic number).

File name: Supplementary Data 11
Description: Information of samples and the survival rates of worker and larvae cared for by Sym-workers and Apo-workers in a 30-day feeding experiment.

File name: Supplementary Data 12
Description: Information of samples and the growth rate of larvae (indicated by larval body length) cared for by Sym-workers and Apo-workers in a 30-day feeding experiment.

### Supplementary Information

#### Symbiotic solutions for colony nutrition: conserved nitrogen recycling within the bacterial pouch of *Tetraponera* ants

Mingjie Ma, Biru Zhu, Dayong Zhang, Piotr Łukasik, Yi Hu

### Supplementary Figures

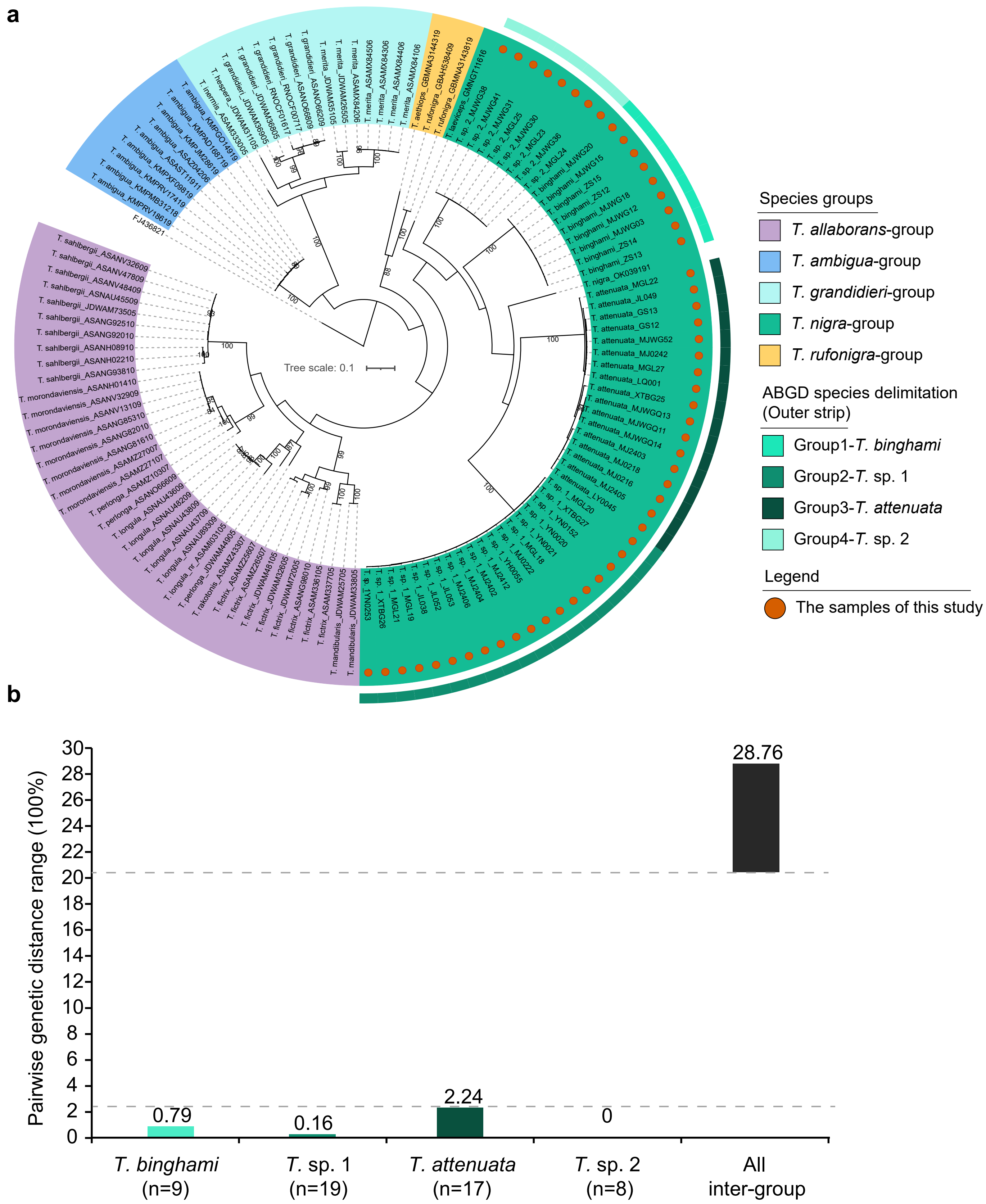

**Supplementary Figure 1. Molecular identification of collected *T. nigra*-group ants. (a)** Rooted maximum likelihood phylogeny inferred from COI sequences of *Tetraponera* ants collected in this study alongside public sequences (>600 bp) retrieved from NCBI and BOLD. Inner-circle shadings denote species groups of *Tetraponera* ants, and outer-circle color strips indicate the results of species delimitation analysis based on ABGD (Automatic Barcode Gap Discovery). Orange dots represent the samples used in this work. **(b)** Frequency histogram of K2P (Kimura two-parameter) pairwise genetic distances among COI sequences of collected *T. nigra*-group ants. Combined phylogenetic inference and species-delimitation analyses support the presence of four distinct *T. nigra*-group species.

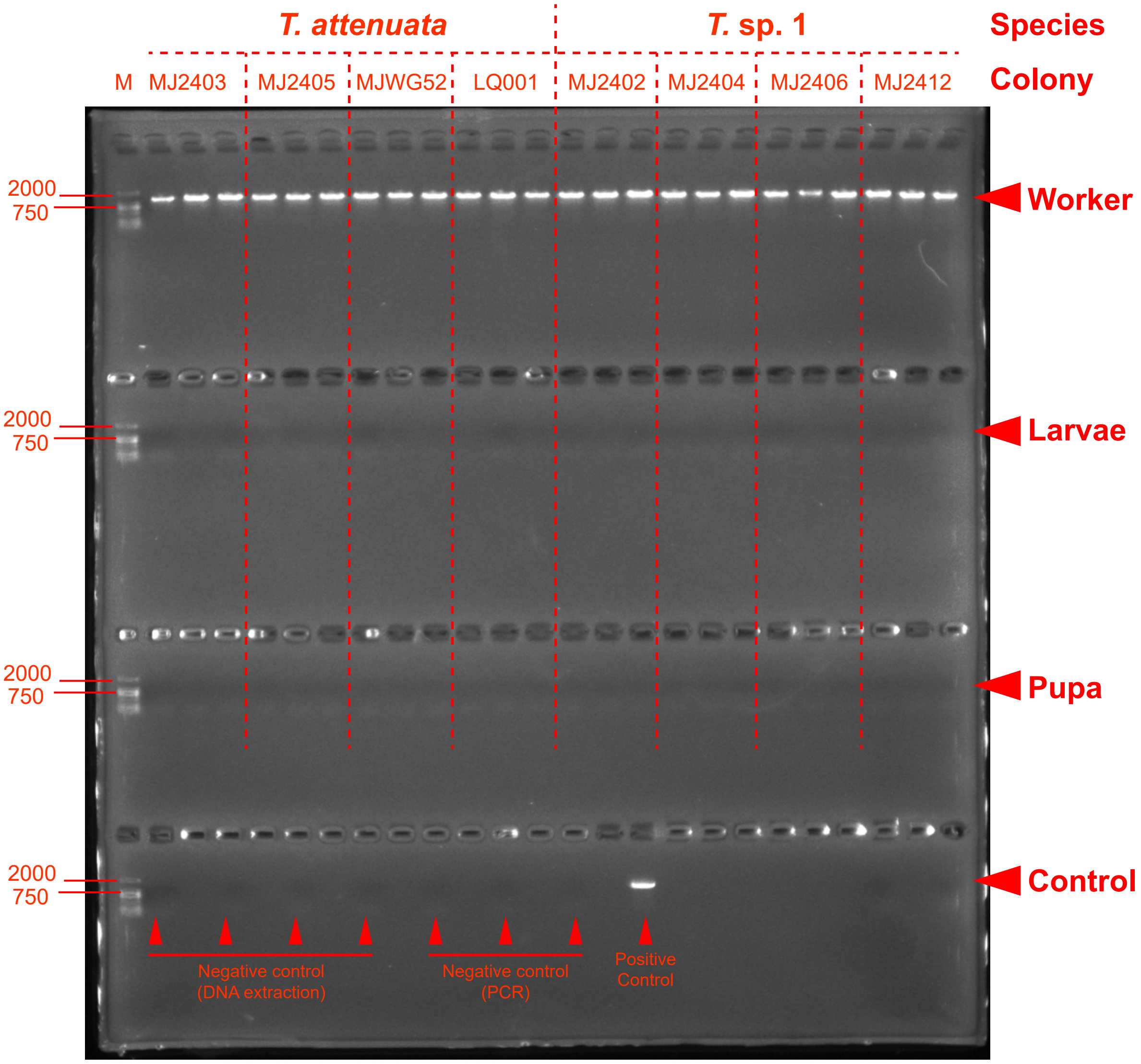

**Supplementary Figure 2. Diagnostic PCR detection of *Tokpelaia* in workers, larvae, and pupae of *T. attenuata* and *T.* sp. 1.** DNA was extracted from the abdomens of workers and whole larvae and pupae. PCR using *Tokpelaia*-specific primers, followed by agarose gel electrophoresis, revealed that *Tokpelaia* was exclusively present in the guts of worker ants, with no detectable presence in larvae or pupae. A 2,000 bp DNA marker was used for size estimation during gel electrophoresis.

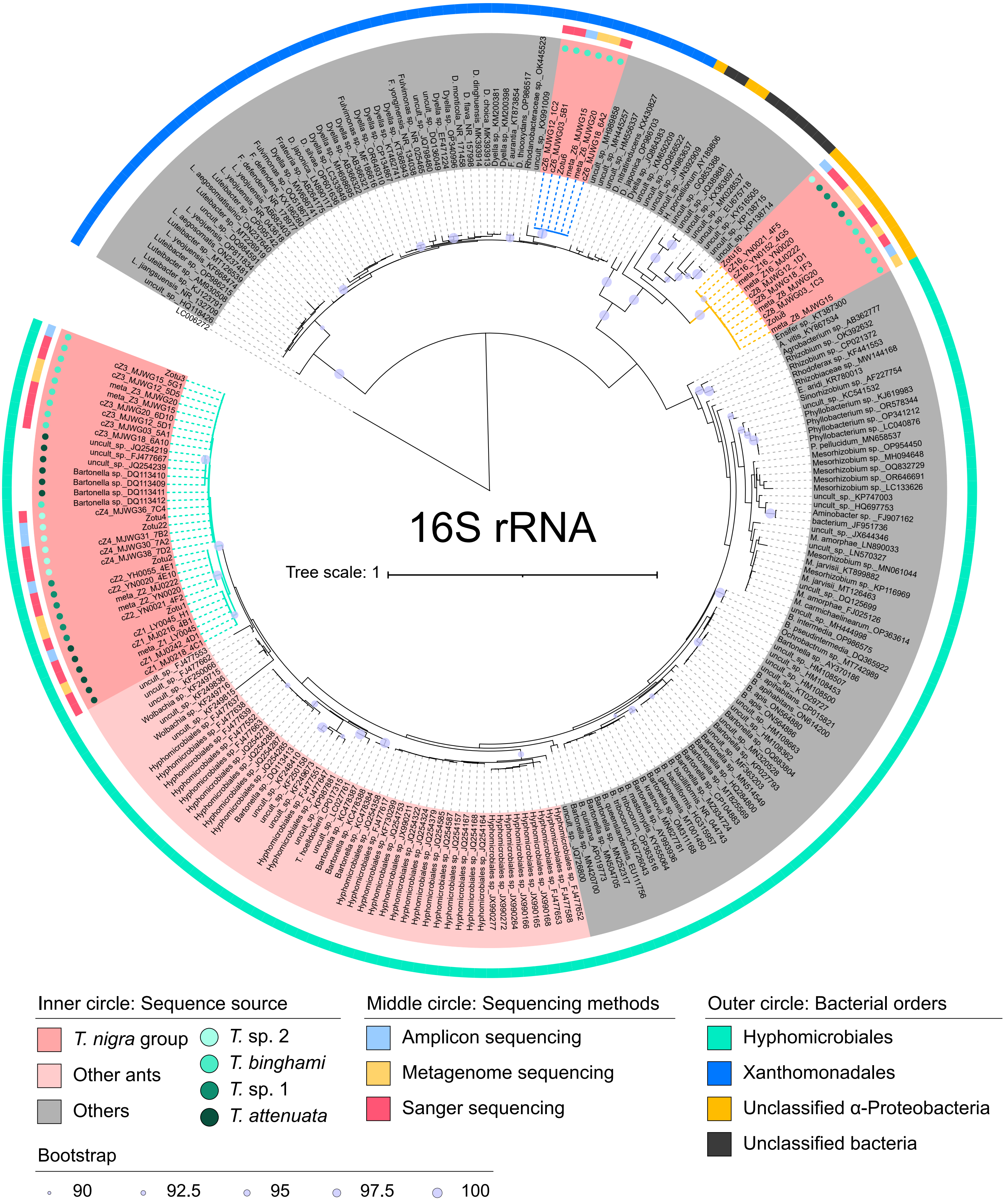

**Supplementary Figure 3. Maximum likelihood phylogeny of 16S rRNA gene sequences demonstrating taxonomic conservation of symbiotic bacteria associated with *Tetraponera* ants.** Sequences were derived from high-abundance zOTUs (>1% relative abundance based on amplicon sequencing datasets), metagenomic data, cloning experiments, and top BLAST hits. The tree reveals that all *Tetraponera*-associated symbionts form host-specific clades. Inner circle: taxon names are color-coded by host origin (red: *T. nigra*-group ants; light red: other ants; gray: non-ant hosts/environments). Middle circle: sequence sources are indicated by colored rings (blue: amplicon sequencing; orange: metagenomics; red: Sanger sequencing). Outer circle: bacterial taxonomy.

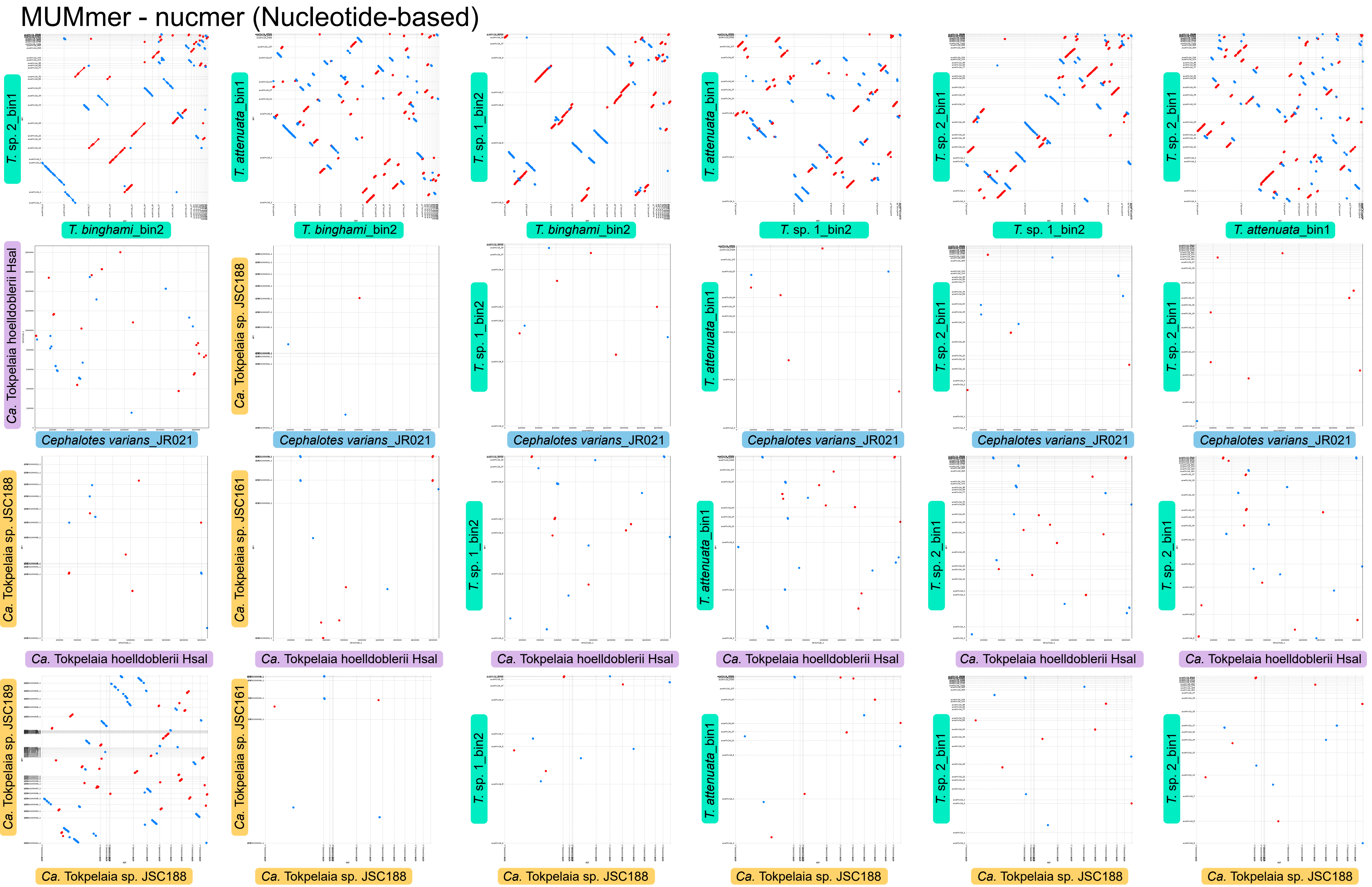

**Supplementary Figure 4. Dot plots of nucleotide-based alignments among different *Tokpelaia* strains.** Comparison of nucleotide order in the genome contigs of *Tokpelaia* symbionts associated with *T. nigra*-group ants and the genomes of *Cephalotes* symbionts, *Ca.* T. hoelldobblerii, and *Dolichoderus* symbionts. Each dot represents a match between nucleotide order. Forward matches are colored in red and reverse matches in blue.

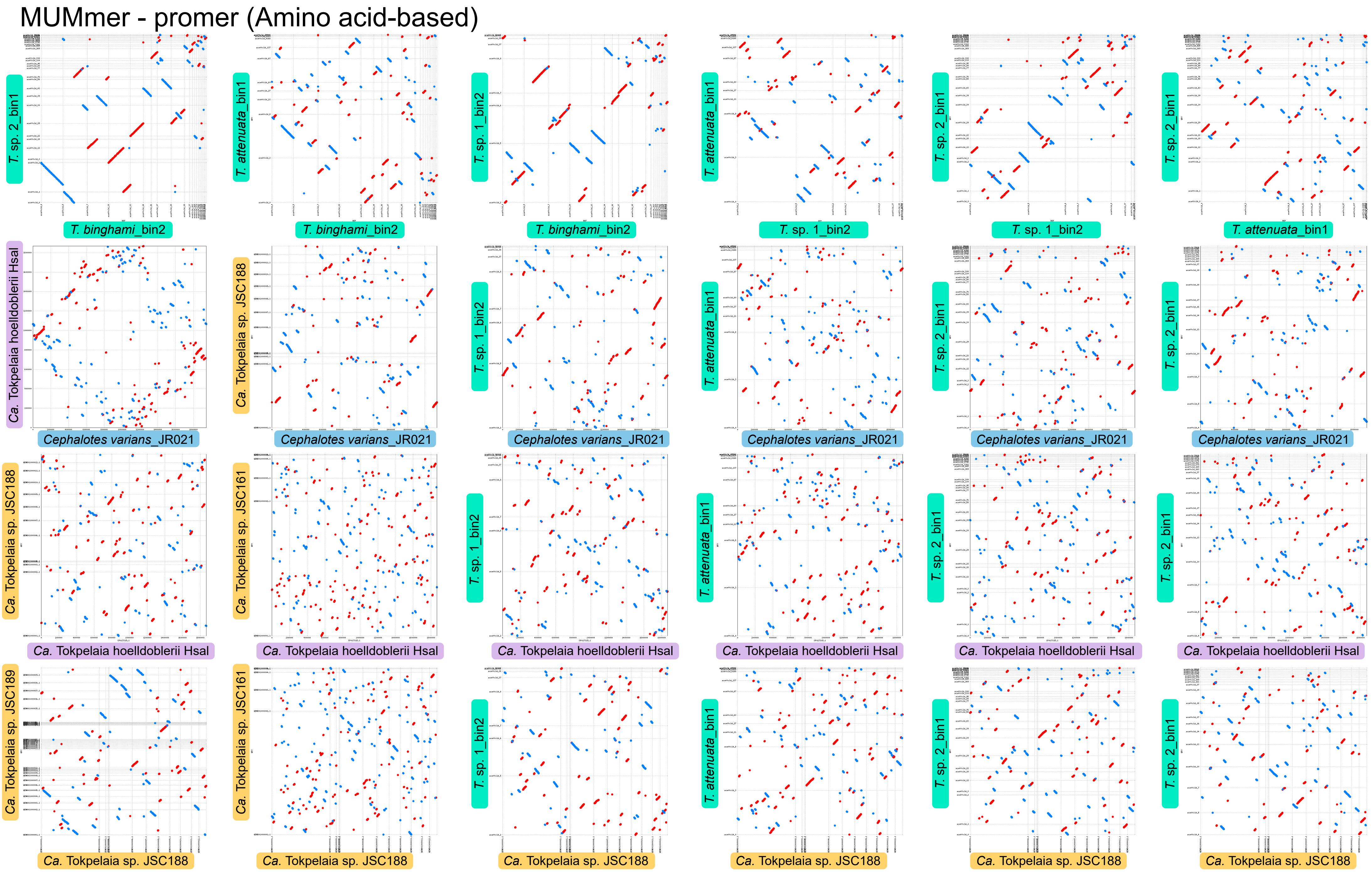

**Supplementary Figure 5. Dot plots of protein-based alignments among different *Tokpelaia* strains.** Comparison of gene order in the genome contigs of *Tokpelaia* symbionts associated with *T. nigra*-group ants and the genomes of *Cephalotes* symbionts, *Ca*. T. hoelldobblerii, and *Dolichoderus* symbionts. Each dot represents a match between Promer-predicted proteins. Forward matches are colored in red and reverse matches in blue.

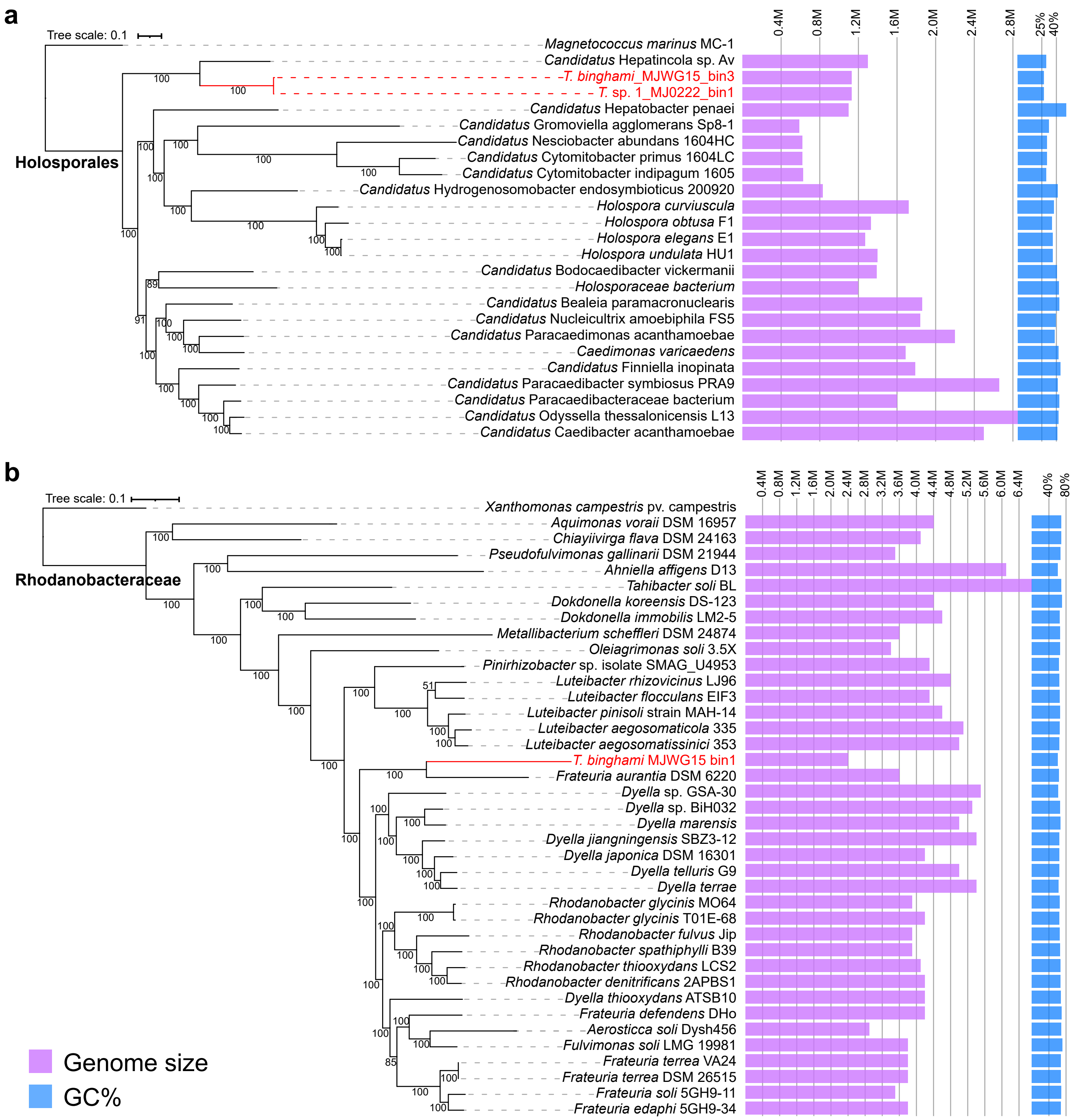

**Supplementary Figure 6. Phylogenomic analysis of unclassified α-Proteobacteria (a) and Xanthomonadales (b) symbionts associated with *T. nigra*-group ants.** Maximum likelihood analysis was performed based on homologous single-copy genes, with genomes from this study highlighted in red. Purple and blue bars represent genome size and GC content, respectively.

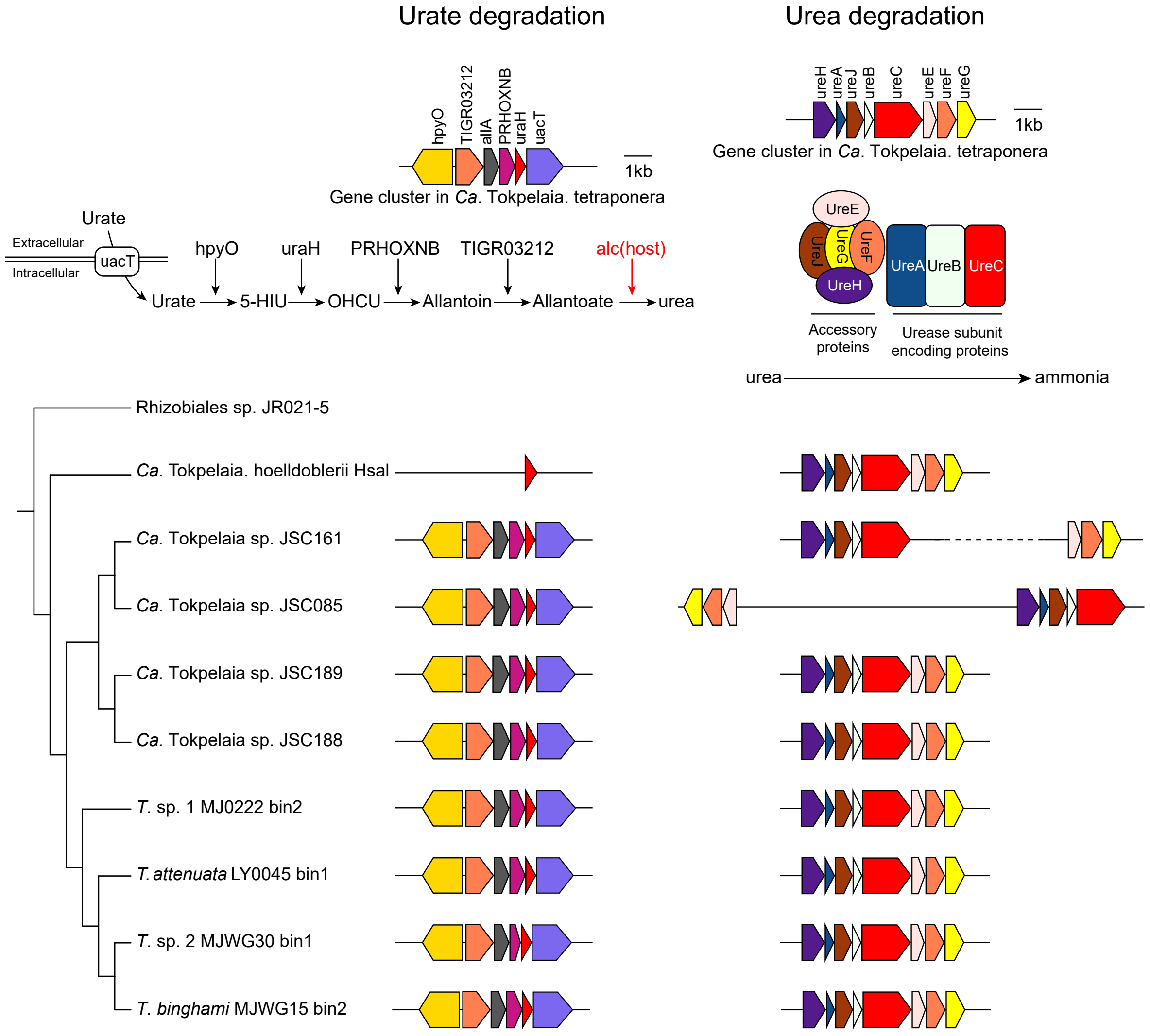

**Supplementary Figure 7. Conserved clusters of genes involved in uric acid degradation and urea degradation pathways across the *Tokpelaia* symbionts of *Tetraponera* and *Dolichoderus* ants.** A cladogram based on the relationship of *Tokpelaia* is shown on the left. Gene annotations for uric acid and urea degradation operons are given at the top of the figure. Genomic organization of these operons is shown across 10 ant-associated *Tokpelaia* genomes, with uric acid degradation gene operons in the left panel and urea degradation gene operons in the right panel. Note that the missing gene alc is present on Hymenoptera-assigned scaffolds in metagenomes of the *T. nigra*-group ants except *T.* sp. 1. A red arrow indicates the host-derived metabolic step.

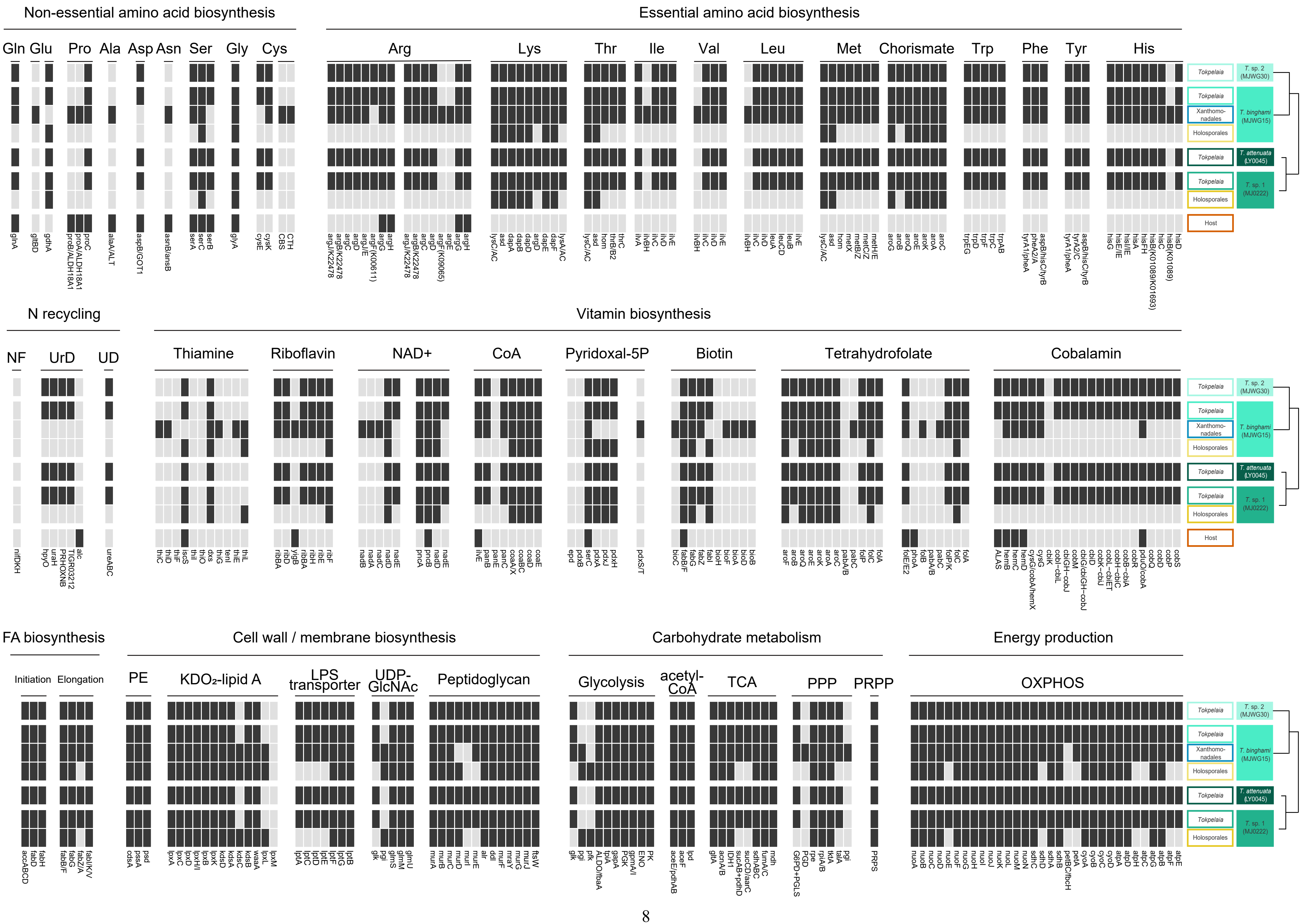
Non-essential amino acid biosynthesis
Essential amino acid biosynthesis
Gln Glu Pro Ala Asp Asn Ser Gly Cys
Arg Lys Thr Ile Val Leu Met Chorismate Trp Phe Tyr His
N recycling
Vitamin biosynthesis
NF UrD UD
Thiamine Riboflavin NAD+ CoA Pyridoxal-5P Biotin Tetrahydrofolate Cobalamin
FA biosynthesis
Cell wall / membrane biosynthesis
Carbohydrate metabolism
Energy production
Initiation Elongation
PE KDO₂-lipid A LPS transporter UDP-GlcNAc Peptidoglycan
Glycolysis acetyl-CoA TCA PPP PRPP
OXPHOS
Tokpelaia
Xanthomonadales
Holosporales
Host
T. sp. 2 (MJWG30)
T. binghami (MJWG15)
T. attenuata (LY0045)
T. sp. 1 (MJ0222)

**Supplementary Figure 8. The comparison of gene sets related to amino acids, vitamins, nitrogen cycling, cell membrane components, fatty acid biosynthesis, carbohydrate metabolism, and energy production from seven symbiont draft genomes.** Each bar represents a single gene with an abbreviation at the bottom. Genes are classified as functional genes (black) and absent (light gray). The maximum likelihood tree based on metagenomic data for four host species is shown on the right. NF: Nitrogen fixation, UrD: Uric acid degradation, UD: Urea degradation, FA: Fatty acid, PE: Phosphatidylethanolamine, PPP: Pentose phosphate pathway, OXPHOS: Oxidative phosphorylation.

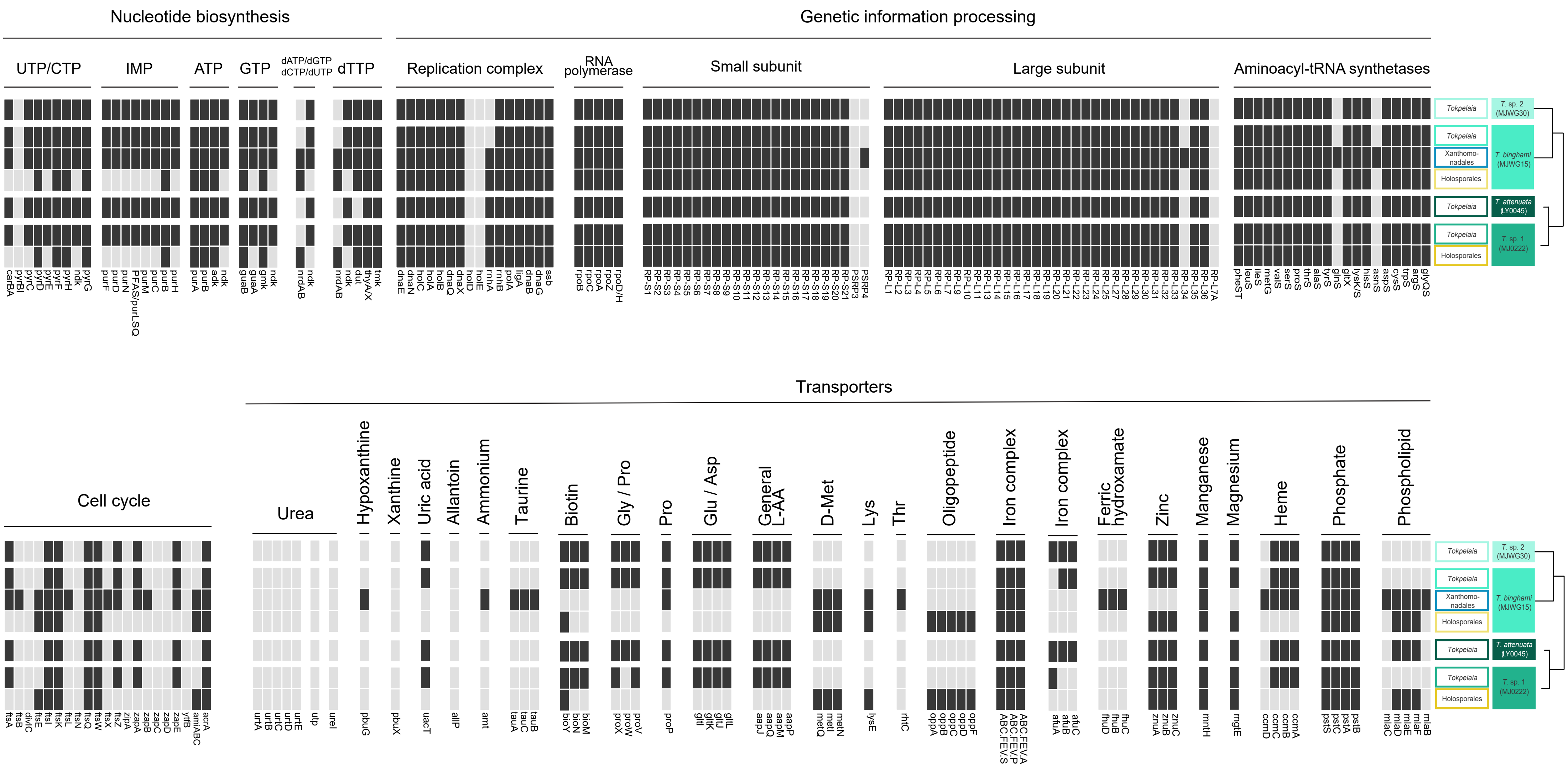

**Supplementary Figure 9. The comparison of gene sets related to nucleotide biosynthesis, genetic information processing, cell cycle, and transporters from seven symbiont draft genomes.** Each bar represents a single gene with an abbreviation at the bottom. Genes are classified as functional genes (black) and absent (light gray). The maximum likelihood tree based on metagenomic data for four host species is shown on the right.

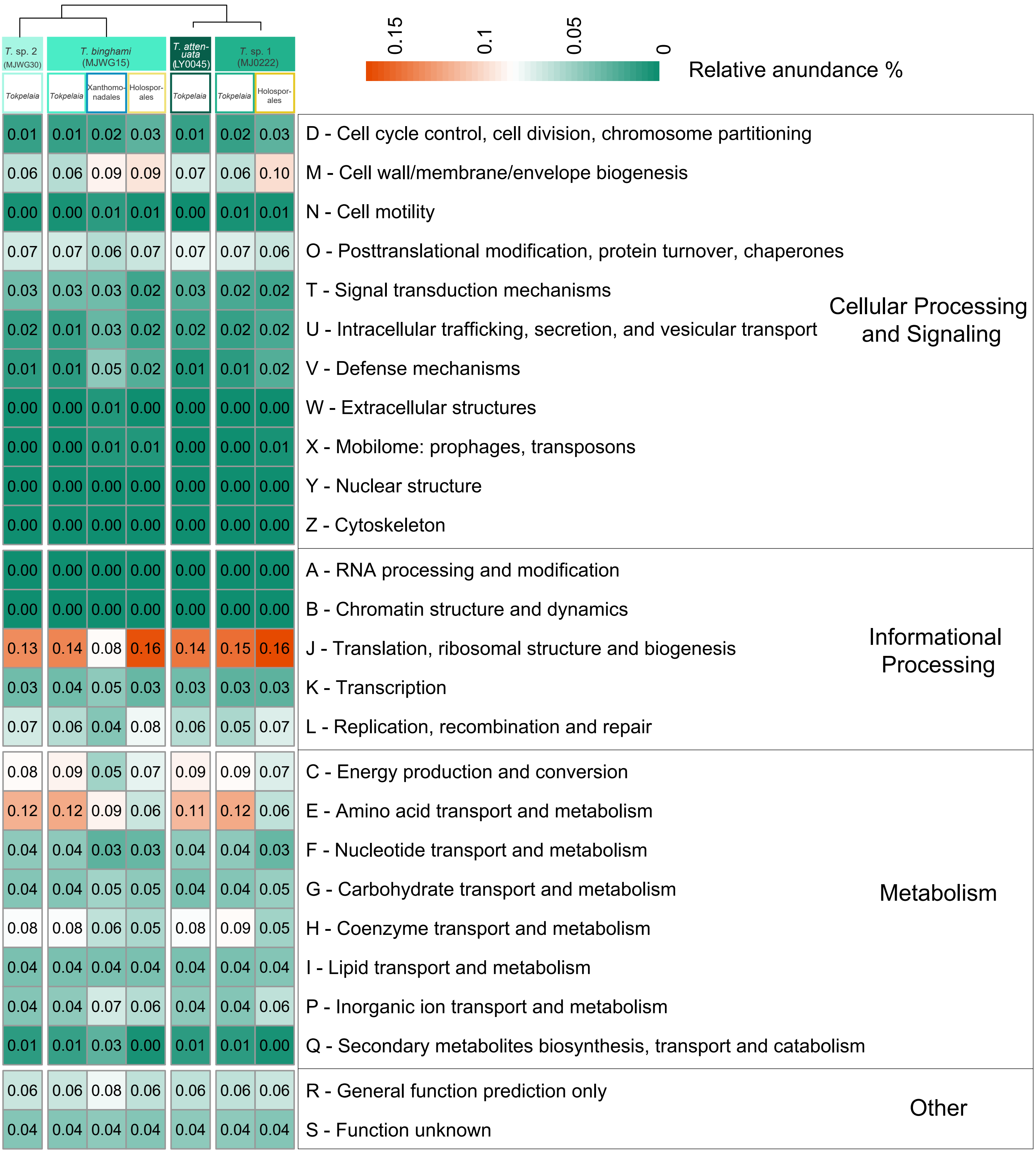

**Supplementary Figure 10. Functional annotation profile of *Tetraponera*-associated symbiont genomes.** Heatmap displays the relative abundance of Clusters of Orthologous Groups (COG) categories (A-Z) across seven symbiont genomes, with color intensity and numeric values representing gene proportions in each functional category.

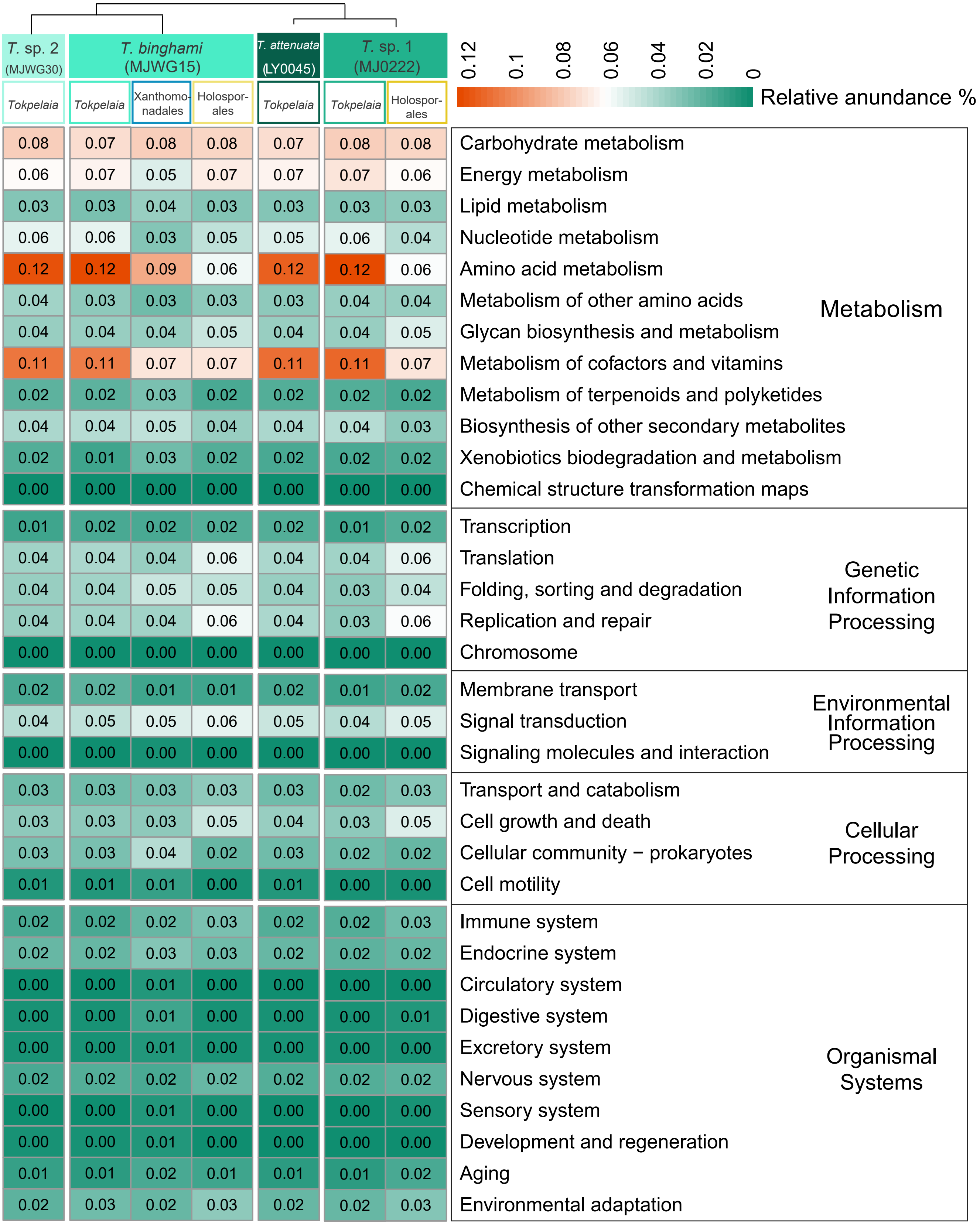

**Supplementary Figure 11. Functional annotation profile of *Tetraponera*-associated symbiont genomes.** Heatmap displays the relative abundance of KEGG functional categories across seven symbiont genomes, with color intensity and numeric values representing gene proportions in each functional category.

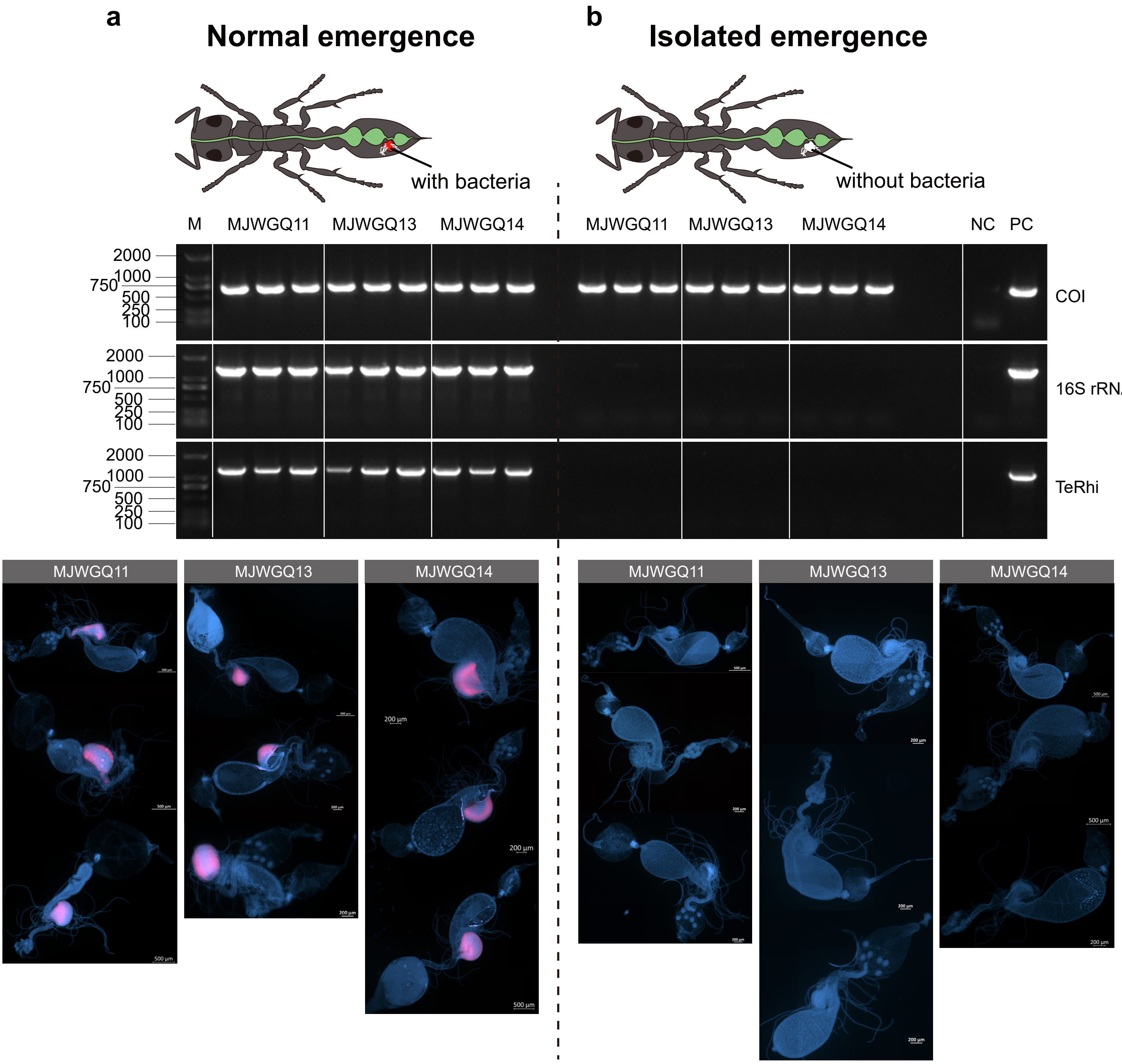

**Supplementary Figure 12. Absence of symbiotic bacteria in bacterial pouches of newly emerged workers reared in isolation.** Three ant colonies (MJWGQ11, MJWGQ13, and MJWGQ14) were selected, and the worker ants from each colony were divided into two groups. Panel a shows workers that emerged under normal conditions, while Panel b represents workers that emerged in isolation, without exposure to nest mates. To confirm the quality of the extracted DNA, we first used the COI gene primers HCO2198 and LCO1490, which produced clear amplification bands for all samples. Subsequently, using universal 16S rRNA primers and *Tetraponera*-specific *Tokpelaia* primers, we found no amplification signals in the DNA of isolated workers from any of the three colonies, while DNA from normally emerged workers consistently produced clear bands. Additionally, using the universal bacterial probe EUB338, no probe signals were detected in the bacterial pouches of isolated workers, while strong signals were observed in the bacterial pouches of normally emerged workers. The fluorescence in situ hybridization (FISH) results show bacterial signals (EUB338) in red and DNA (DAPI) in blue, confirming the presence of bacteria only in normally emerged workers.

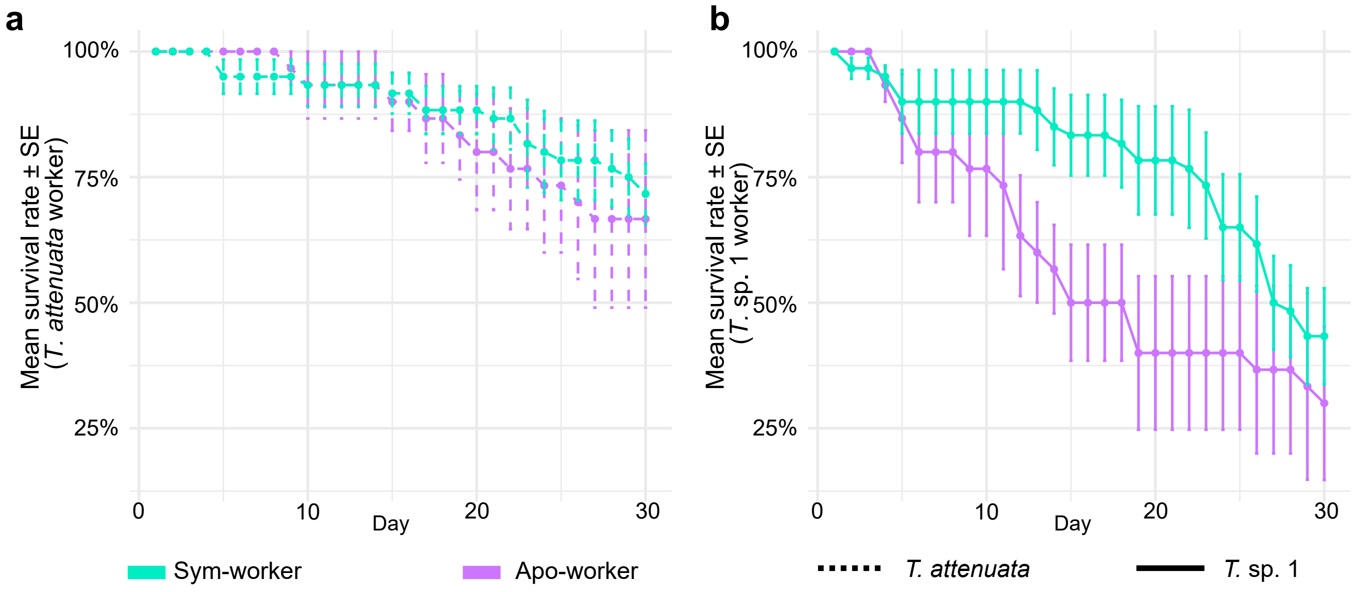

**Supplementary Figure 13.** Survival rates of Apo-workers and Sym-workers of *T. attenuata* (a) and *T.* sp. 1 (b) over a 30-day experimental period.

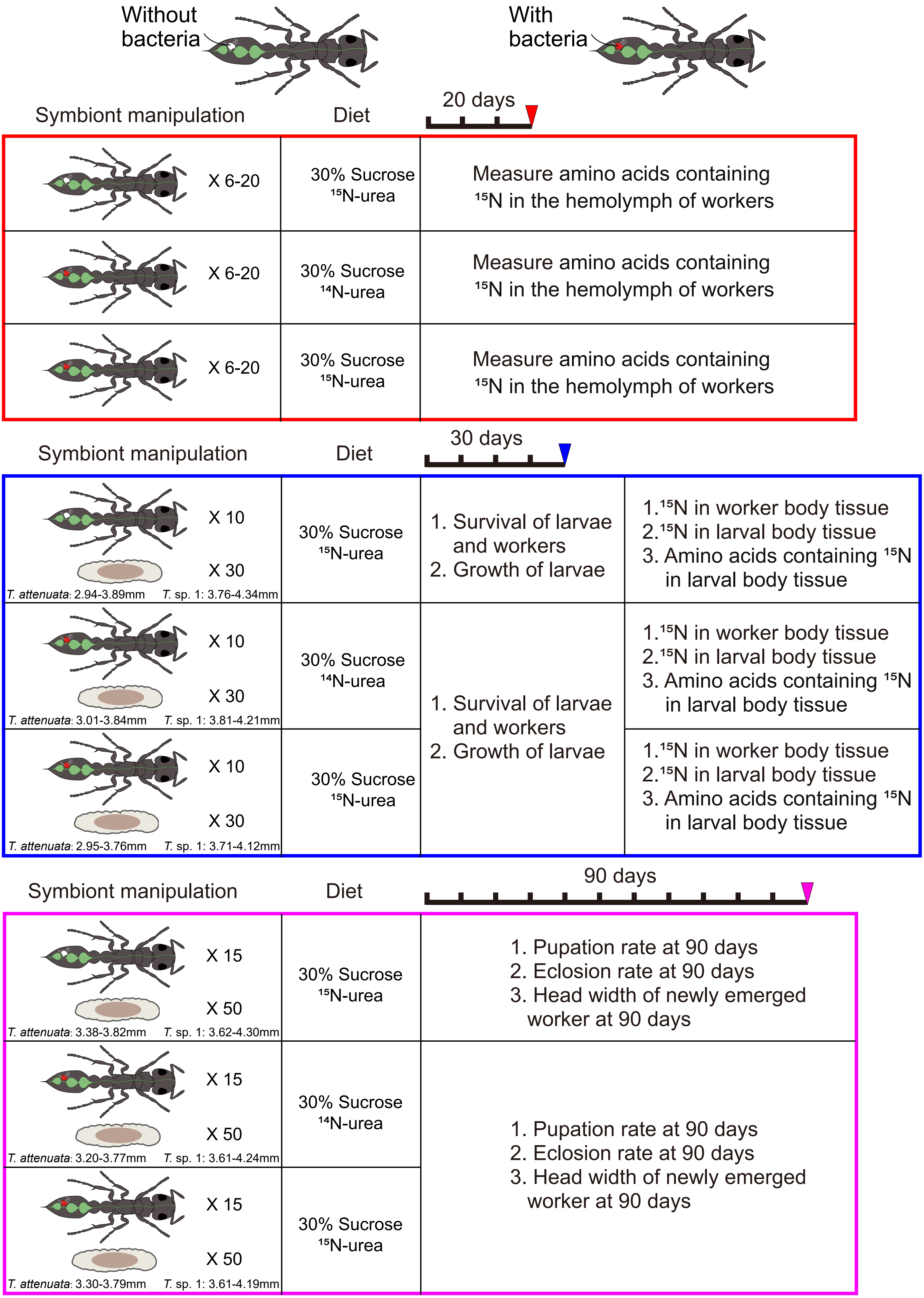

**Supplementary Figure 14. Schematic overview of feeding experiments.** The three colored boxes represent three independent experiments. Each box summarizes the experimental design, fitness parameters recorded throughout the experiment, and metabolomic and isotopic profiling performed at the end of the trial. The initial body length range of larvae used in each experiment is indicated below the larval illustration.

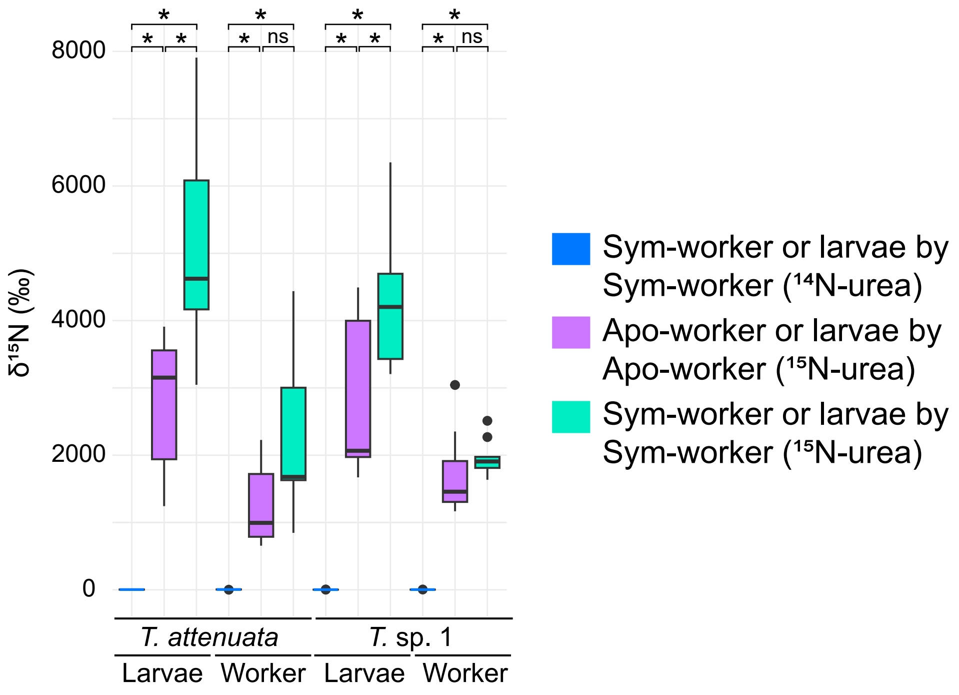

**Supplementary Figure 15. Isotopic enrichment analysis of larval and worker tissues under different symbiotic conditions.** $\delta^{15}N$ values (mean ± SD) are shown for whole-body tissues of gut-resected larvae and abdomen-excised workers in symbiont-containing (sym-worker) and symbiont-free (apo-worker) treatment groups.

### Supplementary Tables

#### Supplementary Table 1. Binning information of metagenomic data.

| Species | Bin ID | Bacterial order | total length | scaffold numbers | N50 [bp] | GC% | | Read depth | | complete% | contamination% | CDS | rRNA | tRNA | Pseudogenes | gtdbtk |
|---|---|---|---|---|---|---|---|---|---|---|---|---|---|---|---|---|
| | | | | | | average | SD | average | SD | | | | | | | |
| *T. attenuata* | LY0045_bin1 | Hyphomicrobiales | 1216848 | 12 | 317650 | 48.44 | 0.006 | 2239.5 | 695.05 | 91.05 | 0 | 1066 | 3 | 39 | 79 | d_Bacteria;p_Proteobacteria;c_Alphaproteobacteria;o_Rhizobiales_A;f_Rhizobiaceae_A;g_;s_ |
| *T. binghami* | MJWG15_bin1 | Xanthomonadales | 2359447 | 5 | 1394701 | 60.99 | 0.004 | 140.3 | 38.72 | 92.05 | 0.69 | 2636 | 3 | 47 | 487 | d_Bacteria;p_Proteobacteria;c_Gammaproteobacteria;o_Xanthomonadales;f_Rhodanobacteraceae;g_Frateuria;s_ |
| | MJWG15_bin2 | Hyphomicrobiales | 1228512 | 20 | 156622 | 49.19 | 0.021 | 2114.9 | 663.73 | 91.7 | 0 | 1202 | 3 | 39 | 147 | d_Bacteria;p_Proteobacteria;c_Alphaproteobacteria;o_Rhizobiales_A;f_Rhizobiaceae_A;g_;s_ |
| | MJWG15_bin3 | unclassified α-proteobacteria | 1129743 | 13 | 162904 | 27.4 | 0.013 | 20.0 | 5.99 | 97.8 | 0 | 1071 | 0 | 32 | 106 | d_Bacteria;p_Proteobacteria;c_Alphaproteobacteria;o_WRAU01;f_WRAU01;g_;s_ |
| *T.* sp. 1 | MJ0222_bin1 | unclassified α-proteobacteria | 1133687 | 6 | 282145 | 27.09 | 0.014 | 81.0 | 5.70 | 97.8 | 0 | 1037 | 0 | 32 | 76 | d_Bacteria;p_Proteobacteria;c_Alphaproteobacteria;o_WRAU01;f_WRAU01;g_;s_ |
| | MJ0222_bin2 | Hyphomicrobiales | 1177748 | 9 | 264414 | 47.51 | 0.013 | 4074.4 | 1115.35 | 89.25 | 0.43 | 1037 | 3 | 37 | 89 | d_Bacteria;p_Proteobacteria;c_Alphaproteobacteria;o_Rhizobiales_A;f_Rhizobiaceae_A;g_;s_ |
| *T.* sp. 2 | MJWG30_bin1 | Hyphomicrobiales | 1255394 | 30 | 73118 | 49.53 | 0.013 | 2502.2 | 561.56 | 91.92 | 0 | 1230 | 3 | 40 | 158 | d_Bacteria;p_Proteobacteria;c_Alphaproteobacteria;o_Rhizobiales_A;f_Rhizobiaceae_A;g_;s_ |

**Supplementary Table 2. 16S rRNA identities and the average amino acid identity (AAI) values for the symbionts of *T. nigra*-group *ants* and *Dolichoderus* spp. Symbionts, *Ca*. Tokpelaia hoelldobblerii.**

| | *Ca*. T. hoelldobblerii | *T. attenuata* LY0045_bin1 | *T. bingham*i MJWG15_bin2 | *T*. sp. 1 MJ0222_bin2 | *T*. sp. 2 MJWG30_bin1 | *Dolichoderus* sp. *Ca.* Tokpelaia sp. JSC161 | *Dolichoderus* sp. *Ca.* Tokpelaia sp. JSC188 | *Dolichoderus* sp. *Ca.* Tokpelaia sp. JSC189 |
|---|---|---|---|---|---|---|---|---|
| *T. attenuata* LY0045_bin1 | 95.87/68.38 | | | | | | | |
| *T. bingham*i MJWG15_bin2 | 95.66/68.14 | 97.87/88.77 | | | | | | |
| *T*. sp. 1 MJ0222_bin2 | 95.17/68.12 | 97.23/86.36 | 97.02/87.29 | | | | | |
| *T*. sp. 2 MJWG30_bin1 | 96.02/68.46 | 98.22/89.08 | 99.50/97.23 | 97.51/87.88 | | | | |
| *Dolichoderus* sp. *Ca.* Tokpelaia sp. JSC161 | 95.53/65.44 | 94.61/69.11 | 95.04/69.21 | 95.11/68.64 | 95.53/69.37 | | | |
| *Dolichoderus* sp. *Ca.* Tokpelaia sp. JSC188 | 95.67/65.66 | 96.17/68.99 | 95.82/69.24 | 95.67/69.08 | 96.31/69.13 | 95.25/67.31 | | |
| *Dolichoderus* sp. *Ca.* Tokpelaia sp. JSC189 | 96.51/67.73 | 96.23/71.69 | 96.30/71.76 | 95.81/71.30 | 96.66/71.75 | 95.18/69.08 | 97.02/78.15 | |
| *Dolichoderus* sp. *Ca.* Tokpelaia sp. JSC085 | 95.60/62.33 | 95.17/65.02 | 95.32/65.10 | 95.04/64.72 | 95.74/65.26 | 95.61/65.98 | 96.17/64.56 | 95.53/65.58 |

**Supplementary Table 3. The statistical results for comparison of missing larval number, pupation and eclosion rates, and head size of emerged workers between symbiotic (Sym-workers) and microbiota-depleted (Apo-workers) colonies in a 90-day feeding experiment.**

| | Pupae | | New worker | | Head width of new worker | |
|---|---|---|---|---|---|---|
| Species | *T. attenuata* | *T.* sp. 1 | *T. attenuata* | *T.* sp. 1 | *T. attenuata* | *T.* sp. 1 |
| Bartlett's test | 0.0921 | 0.1241 | 0.0676 | 0.0968 | 0.2221 | -- |
| Shapiro-Wilk normality test | 0.2414 | 0.5599 | 0.0708 | **0.0042** | 0.1631 | 0.2604 |
| Wilcoxon-Mann-Whitney test | -- | -- | -- | 0.1999 | -- | 0.1818 |
| T-test | 0.2326 | **0.0096** | 0.2271 | -- | **0.0453** | -- |

**Supplementary Table 4. Statistical results for heavy isotopic signal in hemolymph amino acids over a 20-day period in $^{15}$N-urea feeding experiment.**

| | $^{15}$N-urea feeding experiment | Ala | Arg | Asn | Cys | Glu | Gln | Gly | His | Ile | Leu | Lys | Met | Phe | Pro | Ser | Thr | Trp | Tyr | Val |
|---|---|---|---|---|---|---|---|---|---|---|---|---|---|---|---|---|---|---|---|---|
| | Shapiro-Wilk normality test | 1.53E-06 | 1.15E-05 | 0.0010 | 4.78E-05 | 3.02E-06 | 2.97E-06 | 1.34E-05 | 5.32E-06 | 0.0026 | 0.0002 | 0.0048 | 0.0003 | 0.0005 | 2.31E-06 | 3.31E-06 | 1.64E-06 | 0.6947 | 9.7340 | 8.48E-06 |
| Parametric test | Effect of diet on the $^{15}$N signal of amino acids in different dietary groups revealed by an ANOVA test | NA | NA | NA | NA | NA | NA | NA | NA | NA | NA | NA | NA | NA | NA | NA | NA | **0.0110** | **2.38E-05** | NA |
| | Pairwise comparison between $^{14}$N-urea vs $^{15}$N-urea-isolated emergence worker using post hoc Tukey HSD test | NA | NA | NA | NA | NA | NA | NA | NA | NA | NA | NA | NA | NA | NA | NA | NA | 0.7525 | 0.9953 | NA |
| | Pairwise comparison between $^{15}$N-urea-worker vs $^{15}$N-urea-isolated emergence worker using post hoc Tukey HSD test | NA | NA | NA | NA | NA | NA | NA | NA | NA | NA | NA | NA | NA | NA | NA | NA | 0.0519 | **7.77E-05** | NA |
| | Pairwise comparison between $^{15}$N-urea vs 14N-urea using post hoc Tukey HSD test | NA | NA | NA | NA | NA | NA | NA | NA | NA | NA | NA | NA | NA | NA | NA | NA | **0.0124** | **0.0001** | NA |
| Non-parametric test | Effect of diet on the $^{15}$N signal of amino acids in different dietary groups revealed by a Kruskal-Wallis test | **0.0070** | **0.0017** | **0.0134** | **0.0292** | **0.0103** | **4.08E-05** | 0.0559 | **0.0041** | **0.0069** | **0.0370** | **0.0159** | 0.3120 | **0.0036** | **0.0028** | **0.0010** | 0.0771 | NA | NA | **0.0049** |
| | Pairwise comparison between $^{14}$N-urea vs $^{15}$N-urea-isolated emergence worker using post hoc Dunn's Test | 0.8474 | 0.5944 | 0.8626 | **0.0133** | 0.7061 | **0.0044** | 1.0000 | 0.8174 | 1.0000 | 0.4431 | 0.5224 | 0.5311 | 0.3038 | 1.0000 | 1.0000 | 1.0000 | NA | NA | 1.0000 |
| | Pairwise comparison between $^{15}$N-urea-worker vs $^{15}$N-urea-isolated emergence worker using post hoc Dunn's Test | **0.0042** | **0.0008** | **0.0367** | 0.1390 | **0.0398** | **0.0465** | 0.0629 | **0.0158** | **0.0091** | **0.0158** | **0.0069** | 0.1966 | **0.0013** | **0.0031** | **0.0009** | 0.0502 | NA | NA | **0.0051** |
| | Pairwise comparison between $^{15}$N-urea vs $^{14}$N-urea using post hoc Dunn's Test | **0.0302** | **0.0186** | **0.0091** | 0.4870 | **0.0061** | **1.04E-05** | 0.0531 | **0.0030** | **0.0107** | 0.2269 | 0.1049 | 0.8857 | 0.0760 | **0.0075** | **0.0052** | 0.1305 | NA | NA | **0.0109** |

**Supplementary Table 5. Statistical results of heavy isotope signals in amino acids from whole larvae reared by *T. attenuata and T.* sp. 1 Sym-workers and Apo-workers over a 30-day period in a $^{15}$N-urea feeding experiment.**

| | $^{15}$N-urea feeding experiment | Ala | Arg | Asn | Glu | Gln | Gly | His | Ile | Leu | Lys | Met | Phe | Pro | Ser | Thr | Trp | Tyr | Val |
|---|---|---|---|---|---|---|---|---|---|---|---|---|---|---|---|---|---|---|---|
| | Shapiro-Wilk normality test | 0.0418 | 7.46E-05 | 0.0221 | 0.0132 | 0.0373 | 0.0024 | 7.17E-05 | 0.0049 | 0.119 | 0.0007 | 0.0007 | 0.0005 | 0.0039 | 0.1265 | 0.2365 | 0.0066 | 0.0004 | 0.0017 |
| | Analysis of Deviance Table (Type III tests) | Species: p=0.6363 Diet: p= 0.0007 Species * Diet =0.4470 | Species: p=0.9798 Diet: p= 1.544e-06 Species * Diet =0.9694 | Species: p=0.9606 Diet: p= 4.477e-05 Species * Diet =0.6386 | Species: p=0.9026 Diet: p= 2.153e-05 Species * Diet =0.9070 | Species: p=0.8953 Diet: p= 0.0046 Species * Diet =0.8792 | Species: p=0.7858 Diet: p= 6.635e-05 Species * Diet =0.4361 | Species: p=0.9979 Diet: p= 0.0139 Species * Diet =0.5537 | Species: p=0.8851 Diet: p= 5.168e-06 Species * Diet =0.6603 | Species: p=0.5653 Diet: p= 6.493e-06 Species * Diet =0.4721 | Species: p=0.9820 Diet: p= 5.607e-16 Species * Diet =0.7936 | Species: p=0.7847 Diet: p= 0.9635 Species * Diet =0.5905 | Species: p=0.3765 Diet: p < 2e-16 Species * Diet =0.4387 | Species: p=0.9118 Diet: p= 0.0007 Species * Diet =0.5928 | Species: p=0.8557 Diet: p= 0.0018 Species * Diet =0.7862 | Species: p=0.9734 Diet: p= 0.4266 Species * Diet =0.8077 | Species: p=0.8822 Diet: p= 0.0005 Species * Diet =0.5209 | Species: p=0.6939 Diet: p= 2.072e-07 Species * Diet =0.8207 | Species: p=0.7889 Diet: p= 0.02986 Species * Diet =0.08134 |
| Parametric test | Effect of diet on the $^{15}$N signal of amino acids in different dietary groups revealed by an ANOVA test | NA | NA | NA | NA | NA | NA | NA | NA | **0.0001** | NA | NA | NA | NA | **0.0007** | 0.0965 | NA | NA | NA |
| | Pairwise comparison between $^{15}$N-urea-Apo-workers vs $^{14}$N-urea-Sym-workers using Tukey HSD test | NA | NA | NA | NA | NA | NA | NA | NA | 0.2182 | NA | NA | NA | NA | 0.0901 | 0.0983 | NA | NA | NA |
| | Pairwise comparison between $^{15}$N-urea-Sym-workers vs $^{15}$N-urea-Apo-workers using Tukey HSD test | NA | NA | NA | NA | NA | NA | NA | NA | **0.0027** | NA | NA | NA | NA | **0.0408** | 0.8758 | NA | NA | NA |
| | Pairwise comparison between $^{15}$N-urea-Sym-workers vs $^{14}$N-urea-Sym-workers using Tukey HSD test | NA | NA | NA | NA | NA | NA | NA | NA | **0.0001** | NA | NA | NA | NA | **0.0005** | 0.2243 | NA | NA | NA |
| Non-parametric test | Effect of diet on the $^{15}$N signal of amino acids in different dietary groups revealed by a Kruskal-Wallis test | **0.0243** | 0.0028 | 0.0017 | 0.0009 | 0.0028 | 0.0031 | **0.0011** | 0.0032 | NA | 0.0017 | 0.1402 | 0.0023 | 0.0011 | NA | NA | 0.0021 | 0.0034 | 0.0021 |
| | Pairwise comparison between $^{15}$N-urea-Apo-workers vs $^{14}$N-urea-Sym-workers using post hoc Dunn's Test | 0.0524 | 0.7746 | **0.0301** | 0.0524 | **0.0142** | 0.2058 | 0.3305 | 1.0000 | NA | 0.3513 | 0.7746 | 0.5804 | **0.0458** | NA | NA | 0.3204 | 1.0000 | 0.4956 |
| | Pairwise comparison between $^{15}$N-urea-Sym-workers vs $^{15}$N-urea-Apo-workers using post hoc Dunn's Test | 0.2164 | **0.0142** | 0.3513 | 0.1571 | 0.7746 | 0.0861 | **0.0249** | **0.0087** | NA | **0.0007** | **0.0774** | **0.0012** | 0.1950 | NA | NA | **0.0399** | **0.0048** | **0.0224** |
| | Pairwise comparison between $^{15}$N-urea-Sym-workers vs $^{14}$N-urea-Sym-workers using post hoc Dunn's Test | **0.0005** | **0.0018** | **0.0007** | **0.0003** | **0.0018** | **0.0011** | **0.0004** | **0.0031** | NA | **0.0301** | 0.2915 | **0.0193** | **0.0004** | NA | NA | **0.0008** | **0.0057** | **0.0009** |

**Supplementary Table 6. Statistical analysis of $^{15}$N relative abundance in the whole bodies of workers and larvae cared for by Sym-workers and Apo-workers over a 30-day period in a $^{15}$N-urea feeding experiment.**

| | | *T. attenuata* | | *T.* sp. 1 | |
|---|---|---|---|---|---|
| | | **larvae** | **worker** | **larvae** | **worker** |
| | Shapiro-Wilk normality test | 1.1360E-02 | 3.6930E-03 | 7.1440E-03 | 2.9640E-03 |
| Non-parametric test | The impact of different treatments on $\delta^{15}$N of larvae and workers revealed by a Kruskal-Wallis test | **1.1360E-02** | **3.6930E-03** | **7.1440E-03** | **2.9640E-03** |
| | Pairwise comparison between $^{15}$N-urea-Apo-worker vs $^{14}$N-urea-Sym-worker using post hoc Dunn's Test | **5.1130E-04** | **2.0380E-03** | **5.6840E-04** | **1.0150E-03** |
| | Pairwise comparison between $^{15}$N-urea-Sym-worker vs $^{15}$N-urea-Apo-worker using post hoc Dunn's Test | **0.0373** | 0.1062 | **0.0399** | 0.1052 |
| | Pairwise comparison between $^{15}$N-urea-Sym-worker vs $^{14}$N-urea-Sym-worker using post hoc Dunn's Test | **1.5027E-04** | **7.6781E-04** | **1.6976E-04** | **3.2960E-04** |

**Supplementary Table 7. Assembly statistics of metagenomic data.**

| Ant species | Colony | Read numbers | Total length Mbp | Total ≥1 Kbp scaffold length(Mbp) | Scaffold number | Scaffold number (≥1Kbp) | N50 for scaffold length [bp] | Largest Contig | GC% mean | Read coverage mean | IMG project ID |
|---|---|---|---|---|---|---|---|---|---|---|---|
| ***T. attenuata*** | LY0045 | 22065240 | 336.29 | 252.52 | 488009 | 50749 | 7540 | 323061 | 40 | 8.0993 | Gp0646313 |
| ***T. binghami*** | MJWG15 | 19108416 | 276.53 | 217.18 | 326970 | 68659 | 3593 | 1394701 | 39.88 | 9.56093 | Gp0688973 |
| ***T.* sp. 1** | MJ0222 | 22619091 | 280.55 | 201.82 | 381087 | 77311 | 2620 | 535206 | 40.6 | 7.09185 | Gp0646295 |
| ***T.* sp. 2** | MJWG30 | 19285212 | 284.57 | 206.7 | 372476 | 78795 | 2615 | 219025 | 40.35 | 8.1488 | Gp0754593 |

**Supplementary Table 8. PCR details.**

| Gene | Primer 1 | Primer 2 | PCR Cocktail (25 ul volume) | PCR Cycling Conditions |
|---|---|---|---|---|
| COI | LCO1490 (5'-GGTCAACAAATCATAAAGATATTGG-3') | HCO2198（5‘-TAAACTTCAGGGTGACCAAAAAATCA-3’） | 12.5µl of 2X MyTaq™ HS Red Mix (Bioline USA, Inc; Randolph, MA), 1.25 µl of each primer at 5 µM (Sangon Biotech, Shanghai, Co., Ltd.), 6.5µl of molecular water (Sigma) and 1µl of DNA template | 1 cycle of 95°C for 120s; 15 cycles of 15s at 95°C, 56℃->51.8°C (decreasing by 0.5℃ each cycle) for 15 s and 30s at 72°C; 35 cycles of 15s at 95℃, 46℃ at 15s and 30s at 72°C; 1 cycle of 72°C for 1min |
| 16S rRNA | 9Fa-6FAM (5'-GAGTTTGATCITIGCTCAG -3') | 1513R (5'-TACIGITACCTTGTTACGACTT-3') | 12.5µl of 2X MyTaq™ HS Red Mix (Bioline USA, Inc; Randolph, MA), 1.25 µl of each primer at 5 µM (Sangon Biotech, Shanghai, Co., Ltd.), 6.5µl of molecular water (Sigma) and 1µl of DNA template | 1 cycle of 95°C for 1min; 40 cycles of 15s at 95°C, 15s at 55°C and 20s at 72°C; 1 cycle of 72°C for 2min |
| 16S rRNA of *Tetraponera*-specific *Tokpelaia* | TeRhiF2 (5'-AGWACTAYGGRATAACACAGAGAAATT -3') | TeRhiR (5'-TCCTTGMGGTTARCACAGCACC-3') | 12.5µl of 2X MyTaq™ HS Red Mix (Bioline USA, Inc; Randolph, MA), 1.25 µl of each primer at 5 µM (Sangon Biotech, Shanghai, Co., Ltd.), 6.5µl of molecular water (Sigma) and 1µl of DNA template | 1 cycle of 95°C for 1min; 35 cycles of 15s at 95°C, 15s at 56°C and 20s at 72°C; 1 cycle of 72°C for 2min |
| 16S rRNA of T vector | M13F (5'-GTAAAACGACGGCCAGT-3') | M13R (5'-GGAAACAGCATAGACCAT-3') | 12.5µl of 2X MyTaq™ HS Red Mix (Bioline USA, Inc; Randolph, MA), 1.25 µl of each primer at 5 µM (Sangon Biotech, Shanghai, Co., Ltd.), 6.5µl of molecular water (Sigma) and 1µl of DNA template | 1 cycle of 95°C for 1min; 40 cycles of 15s at 95°C, 15s at 55°C and 30s at 72°C; 1 cycle of 72°C for 2min |

**Supplementary Table 9. The number of pupae, new workers cared for by Sym-workers and Apo-workers within a 90-day period.**

| Groups | Species | Colony | Diet | With or without bacteria | Total number of larvae | # Pupae | # New worker |
|---|---|---|---|---|---|---|---|
| MJ2402_N14 | *T.* sp. 1 | MJ2402 | 14N-urea | Sym-worker | 50 | 7 | 2 |
| MJ2402_DN15 | *T.* sp. 1 | MJ2402 | 15N-urea | Apo-worker | 50 | 0 | 0 |
| MJ2402_N15 | *T.* sp. 1 | MJ2402 | 15N-urea | Sym-worker | 50 | 2 | 1 |
| MJ2404_N14 | *T.* sp. 1 | MJ2404 | 14N-urea | Sym-worker | 50 | 4 | 0 |
| MJ2404_DN15 | *T.* sp. 1 | MJ2404 | 15N-urea | Apo-worker | 50 | 1 | 0 |
| MJ2404_N15 | *T.* sp. 1 | MJ2404 | 15N-urea | Sym-worker | 50 | 5 | 1 |
| MJ2406_N14 | *T.* sp. 1 | MJ2406 | 14N-urea | Sym-worker | 50 | 8 | 4 |
| MJ2406_DN15 | *T.* sp. 1 | MJ2406 | 15N-urea | Apo-worker | 50 | 2 | 0 |
| MJ2406_N15 | *T.* sp. 1 | MJ2406 | 15N-urea | Sym-worker | 50 | 3 | 0 |
| MJ2412_N14 | *T.* sp. 1 | MJ2412 | 14N-urea | Sym-worker | 50 | 3 | 2 |
| MJ2412_DN15 | *T.* sp. 1 | MJ2412 | 15N-urea | Apo-worker | 50 | 1 | 1 |
| MJ2412_N15 | *T.* sp. 1 | MJ2412 | 15N-urea | Sym-worker | 50 | 4 | 0 |
| MJ2403_N14 | *T. attenuata* | MJ2403 | 14N-urea | Sym-worker | 50 | 9 | 4 |
| MJ2403_DN15 | *T. attenuata* | MJ2403 | 15N-urea | Apo-worker | 50 | 4 | 0 |
| MJ2403_N15 | *T. attenuata* | MJ2403 | 15N-urea | Sym-worker | 50 | 6 | 4 |
| MJ2405_N14 | *T. attenuata* | MJ2405 | 14N-urea | Sym-worker | 50 | 11 | 2 |
| MJ2405_DN15 | *T. attenuata* | MJ2405 | 15N-urea | Apo-worker | 50 | 1 | 1 |
| MJ2405_N15 | *T. attenuata* | MJ2405 | 15N-urea | Sym-worker | 50 | 16 | 7 |
| MJ2413_N14 | *T. attenuata* | MJ2413 | 14N-urea | Sym-worker | 50 | 0 | 0 |
| MJ2413_DN15 | *T. attenuata* | MJ2413 | 15N-urea | Apo-worker | 50 | 2 | 1 |
| MJ2413_N15 | *T. attenuata* | MJ2413 | 15N-urea | Sym-worker | 50 | 1 | 0 |

**Supplementary Table 10. The head width of worker that have eclosed from larvae cared for by Sym-workers and Apo-workers over a 90-day period.**

| Sample ID | Species | Colony | Diet | With or without bacteria | Width (mm) |
|---|---|---|---|---|---|
| MJ24_03_DN15_1 | *T. attenuata* | MJ2403 | 15N-urea | Apo-worker | 1.105 |
| MJ24_05_DN15_1 | *T. attenuata* | MJ2405 | 15N-urea | Apo-worker | 1.121 |
| MJ24_05_N14_1 | *T. attenuata* | MJ2405 | 14N-urea | Sym-worker | 1.149 |
| MJ24_05_N14_2 | *T. attenuata* | MJ2405 | 14N-urea | Sym-worker | 1.233 |
| MJ24_05_N15_1 | *T. attenuata* | MJ2405 | 15N-urea | Sym-worker | 1.137 |
| MJ24_05_N15_2 | *T. attenuata* | MJ2405 | 15N-urea | Sym-worker | 1.217 |
| MJ24_05_N15_3 | *T. attenuata* | MJ2405 | 15N-urea | Sym-worker | 1.156 |
| MJ24_05_N15_4 | *T. attenuata* | MJ2405 | 15N-urea | Sym-worker | 1.234 |
| MJ24_05_N15_5 | *T. attenuata* | MJ2405 | 15N-urea | Sym-worker | 1.219 |
| MJ24_05_N15_6 | *T. attenuata* | MJ2405 | 15N-urea | Sym-worker | 1.227 |
| MJ24_05_N15_7 | *T. attenuata* | MJ2405 | 15N-urea | Sym-worker | 1.268 |
| MJ24_13_N14_1 | *T. attenuata* | MJ2413 | 14N-urea | Sym-worker | 1.123 |
| MJ24_13_N14_2 | *T. attenuata* | MJ2413 | 14N-urea | Sym-worker | 1.246 |
| MJ24_13_N14_3 | *T. attenuata* | MJ2413 | 14N-urea | Sym-worker | 1.221 |
| MJ24_13_N14_4 | *T. attenuata* | MJ2413 | 14N-urea | Sym-worker | 1.207 |
| MJ24_13_N15_1 | *T. attenuata* | MJ2413 | 15N-urea | Sym-worker | 1.089 |
| MJ24_13_N15_2 | *T. attenuata* | MJ2413 | 15N-urea | Sym-worker | 1.148 |
| MJ24_13_N15_3 | *T. attenuata* | MJ2413 | 15N-urea | Sym-worker | 1.204 |
| MJ24_13_N15_4 | *T. attenuata* | MJ2413 | 15N-urea | Sym-worker | 1.173 |
| MJ24_02_N14_1 | *T.* sp. 1 | MJ2402 | 14N-urea | Sym-worker | 1.458 |
| MJ24_02_N14_2 | *T.* sp. 1 | MJ2402 | 14N-urea | Sym-worker | 1.451 |
| MJ24_02_N15_1 | *T.* sp. 1 | MJ2402 | 15N-urea | Sym-worker | 1.394 |
| MJ24_04_N15_1 | *T.* sp. 1 | MJ2404 | 15N-urea | Sym-worker | 1.46 |
| MJ24_06_N14_1 | *T.* sp. 1 | MJ2406 | 14N-urea | Sym-worker | 1.4 |
| MJ24_06_N14_2 | *T.* sp. 1 | MJ2406 | 14N-urea | Sym-worker | 1.455 |
| MJ24_06_N14_3 | *T.* sp. 1 | MJ2406 | 14N-urea | Sym-worker | 1.435 |
| MJ24_06_N14_4 | *T.* sp. 1 | MJ2406 | 14N-urea | Sym-worker | 1.376 |
| MJ24_12_DN15_1 | *T.* sp. 1 | MJ2412 | 15N-urea | Apo-worker | 1.285 |
| MJ24_12_N14_1 | *T.* sp. 1 | MJ2412 | 14N-urea | Sym-worker | 1.366 |
| MJ24_12_N14_2 | *T.* sp. 1 | MJ2412 | 14N-urea | Sym-worker | 1.355 |

**Supplementary Table 11. The statistical results for comparison of pupation and eclosion rates, and head size of emerged workers between symbiotic (Sym-workers) and microbiota-depleted (Apo-workers) colonies receiving $^{15}N/^{14}N$-labeled urea feedings in a 90-day feeding experiment.**

| | | Pupation rate | | Eclosion rate | | Head width of new worker | |
|---|---|---|---|---|---|---|---|
| | Species | *T. attenuata* | *T.* sp. 1 | *T. attenuata* | *T.* sp. 1 | *T. attenuata* | *T.* sp. 1 |
| | Shapiro-Wilk normality test | 0.2414 | 0.5599 | 0.0708 | 0.0042 | 0.1631 | 0.2604 |
| Parametric test | ANOVA test | 0.5070 | 0.0116 | 0.359 | -- | 0.136 | 0.0568 |
| Parametric test | Pairwise comparison between $^{15}N$-urea-Apo-worker vs $^{14}N$-urea-Sym-worker using post hoc Tukey HSD test | 0.6355 | **0.0092** | 0.7762 | -- | 0.1313 | 0.0572 |
| Parametric test | Pairwise comparison between $^{15}N$-urea-Sym-worker vs $^{15}N$-urea-Apo-worker using post hoc Tukey HSD test | 0.5159 | 0.1313 | 0.3318 | -- | 0.1519 | 0.0651 |
| Parametric test | Pairwise comparison between $^{15}N$-urea-Sym-worker vs $^{14}N$-urea-Sym-worker using post hoc Tukey HSD test | 0.9743 | 0.2461 | 0.6793 | -- | 0.9446 | 0.9008 |
| Non-parametric test | Kruskal-Wallis test | -- | -- | -- | 0.1406 | -- | -- |
| Non-parametric test | Pairwise comparison between $^{15}N$-urea-Apo-worker vs $^{14}N$-urea-Sym-worker using post hoc Dunn's Test | -- | -- | -- | 0.0855 | -- | -- |
| Non-parametric test | Pairwise comparison between $^{15}N$-urea-Sym-worker vs $^{15}N$-urea-Apo-worker using post hoc Dunn's Test | -- | -- | -- | 0.9513 | -- | -- |
| Non-parametric test | Pairwise comparison between $^{15}N$-urea-Sym-worker vs $^{14}N$-urea-Sym-worker using post hoc Dunn's Test | -- | -- | -- | 0.2302 | -- | -- |

### Supplementary Methods

#### Ant collection and specimen identification

We collected 53 *Tetraponera* colonies from southern provinces of China, including Yunnan, Guangdong, and Jiangxi, between 2019 and 2024 (Supplementary Data 1). For molecular analyses, multiple workers from each colony were preserved in anhydrous ethanol and stored at -20°C before DNA extraction. Colonies designated for dietary manipulation experiments or fluorescence in situ hybridization (FISH) were kept alive for laboratory rearing.

For taxonomic identification, one individual from each colony was photographed and identified based on morphological characteristics. Additionally, we selected another individual per colony for DNA barcoding analysis of the mitochondrial cytochrome oxidase I gene (COI) to refine species identification. Our analysis included all available COI fragment sequences of *Tetraponera* genus representatives obtained from the GenBank and BOLD databases (https://www.boldsystems.org/). Afterward, sequences with a length shorter than 600 bp and those lacking clear classification were excluded. Using the COI sequence of *Pseudomyrmex gracilis* (FJ436821.1), a sister genus of *Tetraponera*, as an outgroup, we aligned all sequences with MAFFT v7.520[1] and removed divergent and ambiguously aligned blocks using Gblocks v0.91b[2]. Phylogenetic analyses were conducted using RAxML v8.2.12[3] with 1000 bootstrap replicates under the GTRGAMMA model, and the resulting phylogeny was visualized using iTOL[4]. Species delimitation analysis was further assessed using the Automatic Barcode Gap Discovery (ABGD) method[5], which detects barcode gaps that separate candidate species by assuming no overlap between intra- and interspecific genetic distances. Pairwise K2P distance matrices were calculated from the COI alignment (excluding the outgroup) using MEGA v11.0.13[6]. These matrices were used for ABGD analysis performed online (http://wwwabi.snv.jussieu.fr/public/abgd/), with a relative gap width (X) of 1.5, and results recorded for intraspecific divergence (P) ranging from 0.001 to 0.1 (Supplementary Fig. 1).

#### Trophic levels of *T. nigra*-group ants determined by nitrogen isotope ratios

While collecting *Tetraponera* ants, we also gathered local plants, herbivores, and predatory arthropods. Plant samples were placed in paper bags containing silica gel, while arthropods were preserved in anhydrous ethanol. Upon returning to the lab, all samples were stored at -20°C. Detailed information regarding plant and arthropod collections is provided in Supplementary Data 3. We removed the abdomens from all arthropod samples before isotopic analysis to prevent contamination from residual food in arthropod guts. Plant leaves and arthropod samples were dried using a vacuum freeze dryer (Shunzhi Inc., Shanghai, China). The dried samples were ground and analyzed for $\delta^{15}N$ analysis using a Delta A Advantage isotope ratio mass spectrometer coupled with an EA-HT elemental analyzer (Thermo Fisher Scientific Inc., Bremen, Germany). The nitrogen isotope ratios expressed as $\delta^{15}N$ value were calculated according to the equation: $\delta(‰) = [(Rsample - Rstandard)/Rstandard]*1000$, where Rsample is the $^{15}N/^{14}N$ ratio of the sample and Rstandard is the $^{15}N/^{14}N$ ratio of atmospheric $N_2$ (Rstandard = 0.003676). To conduct comparisons across different locations, we computed the average $\delta^{15}N$ values for plants ($\delta^{15}N$ plant) examined at each specific site. The relative trophic level of ants and arthropods was then determined by subtracting $\delta^{15}N$ plant from $\delta^{15}N$ ant/arthropod at each location. Finally, we adjusted for local variation by subtracting the median $\delta^{15}N$ value of plants at each site from the overall median value across all locations.

#### Structure of bacterial pouch and bacterial localization

To investigate the organization of the bacterial pouch across different life stages in the *T. nigra*-group ants, we examined the gut structure of workers, pupae and larvae of *T. attenuata* using scanning electron microscopy (SEM). Guts were dissected and fixed overnight at 4°C in 2.5% glutaraldehyde. The samples were then dehydrated through a graded ethanol series (30%, 50%, 70%, 85%, 90%, and 95% ethanol for 15 minutes each, followed by three 15-minute washes in 100% ethanol). After dehydration, the samples were dried using a critical point dryer (Leica CPD300), sputter-coated with gold (Hitachi E-1045), and observed under a scanning electron microscope (Hitachi SU8010, Tokyo, Japan).

Fluorescence In Situ Hybridization (FISH) was conducted to localize symbiotic bacteria in workers of four *T. nigra*-group species, following a previously published protocol[7]. Guts were dissected and fixed in 4% formaldehyde in PBS buffer at room temperature for 2 hours. The samples were then immersed in 80% ethanol containing 6% hydrogen peroxide solution at room temperature for 1-4 weeks to reduce autofluorescence of insect tissues. Subsequently, the samples were rehydrated by washing with PBSTx (PBS with 0.3% Triton X-100) for 10 minutes three times, followed by washing with hybridization buffer (30% formamide, 0.01% SDS, 0.9 M NaCl, and 0.02 M Tris-HCl, pH 8.0) for 10 minutes three times. Finally, the samples were incubated in a hybridization buffer containing 100 nM EUB338 universal bacterial probe and 1% DAPI overnight at room temperature. After washing the specimens three times for 10 minutes each with PBSTx, specimens were imaged using an inverted fluorescence microscope (Observer Z1, Zeiss, Germany). To further identify specific symbionts, a second round of FISH was performed using species-specific probes targeting Rhizobiales, including RHIZ1244[8] (Cy5-TCGCTGCCCACTGTCACC) for *T. attenuata* and *T.* sp. 2, TBIRHI (Cy5-TCGCTACCCACTGTCACC) for *T. binghami*, and TSP1RHI (Cy5-TCGCTACCCATTGTCACC) for *T.* sp. 1.

To obtain detailed structure information on the bacterial pouch and associated bacteria, we performed semithin sectioning and transmission electron microscopy (TEM). The guts of *T. attenuata* were dissected and fixed in 2.5% (vol/vol) glutaraldehyde in phosphate buffer saline (PBS) (0.1 M, pH 7.4) at 4°C overnight. After washing with PBS four times, the samples were postfixed with 1% (w/v) osmium tetraoxide in 0.1 M PBS for 2 h at room temperature. Then, the samples were dehydrated in a graded ethanol series (30%, 50%, 70%, 80%, 90% ethanol for 7 minutes once; 100% ethanol for 7 minutes twice) and then in pure acetone for 10 minutes twice. Then, the samples were infiltrated in a graded mixture (3:1, 1:1, 1:3) of acetone and SPI-PON812 resin (16.2 g SPI-PON812, 10 g DDSA and 8.9 g NMA) and then infiltrated in pure resin. Finally, tissues were embedded in pure resin with 1.5% BDMA and polymerized at 45°C for 12h and then at 60°C for 48 h. The ultrathin sections (70nm thick) were sectioned with microtome (Leica EM UC6), double-stained with uranyl acetate and lead citrate, and examined under a transmission electron microscope (Tecnai Spirit120kV, FEI, USA).

#### DNA extraction

Ant workers underwent surface sterilization by immersion in 6% bleach for one minute, followed by a single wash with 95–100% ethanol and three subsequent washes with molecular-grade water. For samples intended for 16S rRNA amplicon sequencing, the gaster of a single worker was dissected using sterile forceps. For metagenomic sequencing, we used sterile forceps to dissect the gut tissues of 10 workers from the same ant colony and combined them into a single sample. Each dissected gaster or pooled gut sample was placed into sterile 1.5 mL tubes containing 180 μL enzymatic lysis buffer, then flash-frozen in liquid nitrogen and ground with sterile pestles. Subsequent extractions were carried out using DNeasy Blood and Tissue kits (Qiagen Ltd., Hilden, Germany) following the manufacturer's protocol for Gram-positive bacteria. To monitor potential contamination from laboratory or reagent sources during DNA extraction, we included blank samples in each extraction batch. DNA quality was assessed by amplifying the COI gene with primers LCO1490 and HCO2198. Details on PCR primers, reaction conditions, and cycling protocols are provided in Supplementary Table 8, with DNA quality data in Supplementary Data 4.

### Bacterial 16S rRNA amplicon sequencing

We investigated the microbial community diversity of *T. nigra*-group ants through the 16S rRNA amplicon sequencing. For each species, samples were collected from 4-6 colonies, with 3-5 workers sampled per colony. We submitted 99 ant samples to Beijing Novogene Bioinformatics Technology Co., Ltd for amplicon library preparation targeting the V4 region of the 16S rRNA gene. Along with insect DNA samples, blank samples in our DNA extraction batches were included to monitor potential contamination during sample preparation and sequencing. In addition, 12 DNA samples from *Escherichia coli* cultures were included to monitor potential cross-contamination during sample preparation and sequencing. Pooled libraries were sequenced on the Illumina NovaSeq6000 SP platform, generating 2x250bp reads (Supplementary Data 4).

The raw reads were demultiplexed, trimmed barcodes and primers, and merged using FLASH v1.2.7[9]. Quality filtering was done with fastp v0.23.4[10], and downstream analysis was performed using a modified version of the "modified_LSD.py" script[11] from https://github.com/catesval/army_ant_myrmecophiles. We made changes to the database in the script to suit our requirements. Briefly, low-quality sequences were removed using VSEARCH v2.15.2[12], followed by chimera removal and denoising with USEARCH v11.0.667[13], which produced zOTU abundance tables. zOTUs were clustered into OTUs based on 97% similarity using the -cluster_otus and -uparseout parameters in usearch, and the resulting clusters were checked for chimeras. Species classification was performed by comparing sequences against the RDP database (http://www.drive5.com/sintax/rdp_16s_v18.fa.gz) using vsearch.

Bacterial DNA present in reagents and the environment can significantly impact microbial community profiles, especially in samples with low template concentrations[14]. To eliminate potential contamination, we implemented several steps following established criteria[7] (Supplementary Data 4). First, we performed BLASTn searches against the NCBI database to assign taxonomy. Sequences identified as mitochondria, chloroplasts, Archaea, Eukarya, or chimera were then removed from further analysis. Next, sequences obtained from cloning amplification experiments (Accession numbers: PQ549644-PQ549668), specifically designed to identify bacterial species in the ant gut, were used as reference sequences. zOTUs that perfectly matched these reference sequences were initially added to the final zOTU list. The remaining zOTUs were then screened for potential contamination. Contaminants were identified by comparing their relative abundance in experimental libraries to that in blank libraries. Specifically, zOTUs with a maximum relative abundance at least ten times greater in any experimental library than in any blank library were considered non-contaminants. Those non-contaminant zOTUs were added to the final zOTU list for further analysis (Supplementary Data 5).

### Metagenomics

For each ant species, DNA was extracted from pooled gut tissues of 10 workers from the same colony and then sent to Beijing Novogene Bioinformatics Technology Co., Ltd for metagenomic sequencing on the Illumina NovaSeq6000 platform. DNA degradation and potential contamination was monitored on 1% agarose gels, and concentrations were measured with a Qubit® dsDNA Assay Kit (Life Technologies, CA, USA). A total amount of 1 μg of DNA per sample was used for library preparation with the NEBNext® Ultra™ DNA Library Prep Kit for Illumina (NEB, USA), following the manufacturer's protocols. DNA fragmentation was performed by sonication to achieve a size of ~350 bp, followed by end repair, A-tailing, and adapter ligation. After PCR amplification, libraries were size-selected using an Agilent 2100 Bioanalyzer and quantified by qPCR. Cluster generation was performed on the cBot system and libraries were sequenced on an Illumina NovaSeq6000 platform to generate paired-end reads.

Quality trimming of raw reads was done with Trimmomatic-0.39[15], followed by read quality checks using FastQC v0.11.9 (http://www.bioinformatics.babraham.ac.uk/projects/fastqc). The trimmed reads were assembled with IDBA-UD 1.1.3[16] using k values of 20, 40, 60, 80, and 100, and assembly statistics were calculated with QUAST-5.0.2[17] (Supplementary Table 7). Scaffolds were uploaded to the Integrated Microbial Genomes with Microbiome Samples Expert Review (IMG/M-ER) website for taxonomic classification and gene annotation[18,19]. We focused on N metabolism by utilizing KEGG as references to manually construct the pathways involved in the degradation of nitrogenous wastes and the biosynthesis of amino acids (Supplementary Data 2).

The bacterial community composition of all metagenomes in this study was visualized by the Taxon-Annotated GC Coverage plots, which employed custom scripts based on a previously published method[20]. Quality-filtered reads were mapped to metagenome scaffolds with BWA 0.7.1[21] using default settings. Scaffold length, GC content, and coverage depth were then calculated using the Perl script sam_len_cov_gc_insert.pl (https://github.com/sujaikumar/assemblage). Taxonomic assignments were then integrated with this data using custom scripts based on a previously published method[20]. The resulting plots simultaneously illustrated three key genomic features, including coverage depth, GC content, length, and taxonomic assignments, enabling a comprehensive assessment of microbiome diversity and dominant symbiont identification (Figure 3).

In order to obtain the draft genomes of individual symbiont strains, we employed the metaWRAP pipeline [22] using CONCOCT, MaxBin, and metaBAT binning modules. The draft genomes were refined with the "Bin_refinement" module, using ≥80% completeness and ≤10% contamination. Few scaffolds belonging to these draft genomes are not classified or not predicted to be present in the *T. nigra*-group ants. To improve accuracy, scaffolds from seven draft genomes were compared against the NCBI database using BLASTn and BLASTx with an e-value threshold of $10^{-6}$ and 70% identity. Scaffolds with unexpected classification or their read coverage significantly deviates from that of most scaffolds within their respective draft genomes were manually removed from the binning datasets (Supplementary Data 6). Bin completeness and contamination were assessed with CheckM[23] and the taxonomic assignment was performed using the Genome Taxonomy Database with GTDB-Tk v2.1.1[24] (Supplementary Table 1).

#### Phylogenetic analyses of symbiotic bacteria and ant hosts

We conducted a phylogenetic analysis to establish the evolutionary relationships among the dominant bacteria in *T. nigra*-group ants based on their 16S rRNA gene sequences. Our dataset included the zOTU sequences with a relative abundance greater than 0.01 in at least one library and present in over 80% of libraries from the same ant species, as well as sequences obtained through clonal amplification and metagenomic sequencing. BLASTn searches were performed to identify their closest relatives. The rationale for including top BLASTn hits was to retain all sequences associated with ants and to select the three most similar sequences from other insects and environmental sources. All sequences were aligned using MAFFT v7.520[1], with divergent regions removed via Gblocks v0.91b[2]. Phylogenetic analysis was conducted with RAxML v8.2.12[3] under the GTRGAMMA model, employing 1000 bootstrap replicates, and visualized using iTOL[4].

We further performed genome-based phylogenetic analysis of symbiotic bacteria in *T. nigra*-group ants. For *Tokpelaia* symbionts, we retrieved all publicly available *Tokpelaia* genomes and incorporated 16 *Bartonella* genomes isolated from bees and mammals. For Xanthomonadales symbionts, 16S rRNA gene phylogenetic analysis revealed that they fell into the subfamily Rhodanobacteraceae clade. Thus, we compiled all available genomes from Rhodanobacteraceae for phylogenetic reconstruction. For unclassified Alphaproteobacteria symbionts, phylogenetic analysis of their 16S rRNA gene sequences provides poor resolution of taxonomic assignments. We thus performed a preliminary classification based on phylogenetic analysis using a representative set of diverse Alphaproteobacteria genomes. This analysis placed the *Tetraponera*-associated symbionts within the Holosporales. Subsequently, we retrieved nearly all available Holosporales genomes for a refined phylogenetic analysis. All genome sequences used for these analyses are listed in Supplementary Data 7. OrthoFinder v2.5.5[25] was used to identify single-copy orthologous genes. Amino acid sequences of these genes were aligned with MAFFT v7.520[1], and divergent/ambiguous regions were removed using Gblocks v0.91b[2]. Alignments were concatenated for phylogenetic inference. ProtTest v3.4.2[26] was used to select the optimal amino acid substitution model. Phylogenetic analysis was then performed with RAxML v8.2.12[3] using the best model and 1000 bootstrap replicates. Results were visualized and annotated on iTOL[4].

In order to clarify the phylogenetic relationships of four *T. nigra*-group ants, we performed phylogenetic analysis of ant hosts using four metagenomic datasets. Single-copy orthologous genes were identified using BUSCO v5.6.1 with its Hymenoptera database[26]. Amino acid sequences were aligned with MAFFT v7.520[1], and divergent regions were removed using Gblocks v0.91b[2]. Alignments were concatenated for phylogenetic inference. The optimal amino acid substitution model was selected using ProtTest v3.4.2[26]. Phylogenetic analysis was conducted using RAxML v8.2.12[3] with 1000 bootstrap replicates, and results were visualized on iTOL[4].

#### Generation of symbiont-free workers via sterile pupal isolation

In order to perform symbiont manipulation experiments, we first obtained large numbers of symbiont-free workers without the use of antibiotics. *T. attenuata* and *T.* sp. 1 colonies were collected from Yunnan Province (Supplementary Data 1) and reared on a diet of 30% honey water and mealworms at 25°C with a 14:10 light-dark cycle. A fresh diet was provided every two days. To generate symbiont-free workers, pupae with pigmented eyes were removed from the original cages and placed in sterile dishes. Those pupae kept under sterile conditions will emerge as adult workers with no bacteria in the bacterial pouch, confirmed by PCR amplification using both universal bacterial (9Fa and 1513R) and *Tokpelaia*-specific 16S rRNA gene primers (TeRhiF2 and TeRhiR), as well as whole-mount FISH staining (Supplementary Figure 12).

#### Feeding experiments with $^{15}N$-labeled urea

Controlled lab experiments were conducted on two ant species, *T. attenuata* and *T.* sp. 1, to quantify the symbionts' contributions to the colony-wide nitrogen economy through N recycling of dietary N waste (Supplementary Figure 14). For each species, three or four colonies were selected as biological replicates. Within each colony, individuals were randomly assigned to one of the three treatments: (1) workers harboring symbiotic bacteria in the bacterial pouch (Sym-worker), fed 30% sucrose water containing 1% (w/v) $^{15}N$-labeled urea; (2) workers without symbiotic bacteria in the bacterial pouch (Apo-worker), fed the same $^{15}N$-labeled urea solution; (3) workers with symbiotic bacteria in the bacterial pouch, fed 30% sucrose water containing 1% (w/v) $^{14}N$-labeled urea as a control. Two feeding experiments were conducted, focusing on the workers and larvae (Supplementary Fig. 14).

We initially conducted the $^{15}N$-labeled urea feeding experiments exclusively with worker ants for 3 weeks. At the end of the experiment, hemolymph was harvested from decapitated ants using borosilicate glass needles. These needles were pulled from microcapillary tubes (1x0.6x100 mm, JITIAN BIO, Beijing) and used to collect droplets exuding from the posterior opening of the head capsule and from the anterior opening of the mesosoma. Based on the number of ants, we aimed to ensure 2–3 replicates for each treatment group within each colony (Supplementary Data 10). Enrichment of amino acids in ant hemolymph was measured at the Scale Biomedicine Technology Co., LTD (Beijing, China). Hemolymph from each replicate was mixed with 1 mL of 80% methanol and processed by ultrasonication. After centrifugation, the supernatant was collected and dried under nitrogen. The residues were then reconstituted in 50μL of 50% acetonitrile for UHPLC-HRMS analysis. A quality control (QC) sample was prepared by pooling equal volumes of all prepared samples. Chromatographic separation was performed using a Thermo Fisher Ultimate 3000 UHPLC system with a Waters BEH Amide column. A 2μl injection was used, with a flow rate of 0.35 mL/min. The mobile phases consisted of water with ammonium acetate (pH 8.5) and acetonitrile/water (90:10, v/v). A gradient elution was applied over 20 minutes. Metabolites were analyzed using a Thermo Fisher Q Exactive™ Hybrid Quadrupole-Orbitrap™ Mass Spectrometer in heated electrospray ionization negative ($HESI^-$) mode. The spray voltage was set to 4000 V, with temperatures set at 320°C for both the capillary and probe heater. The full scan was conducted with a resolution of 70,000 FWHM (m/z = 200) in the range of 180–600 m/z. Mass spectral data were processed using Thermo Fisher Xcalibur software (version 4.0.27.19).

Next, we extended the $^{15}N$-labeled urea feeding experiments to include larvae tended by adult workers, with experimental durations of 30 days and 90 days (Supplementary Fig. 12). In the 30-day experiment, each treatment group was assigned 10 workers and 30 larvae. At the end of the experiment, the $^{15}N$ abundance in larval and worker body tissues, excluding the gut to avoid potential food contamination, was measured using a MAT253 stable isotope mass spectrometer at the Beijing Academy of Agriculture and Forestry Sciences (BAAFS) to determine whether the nitrogen assimilated from urea by symbiotic bacteria was utilized by the host and incorporated into its body tissues. Samples were stored at -20°C for at least 24 hours prior to vacuum freeze-drying. The lyophilized samples were finely ground before analysis. Nitrogen isotope ratios were measured using elemental analyzer–isotope ratio mass spectrometry (EA–IRMS) with a MAT 253 IRMS (Thermo Fisher Scientific, USA) interfaced with a Flash 2000 Elemental Analyzer (Thermo Fisher Scientific, USA). Powdered ant and larval samples (0.4–1.2 mg) were weighed into tin capsules (5 mm × 8 mm), compressed into small squares, and introduced into the elemental analyzer via an auto-sampler. Capsules were combusted at 960°C in a helium atmosphere with a pulse of oxygen and subsequently reduced to $N_2$ gas by copper particles in the elemental analyzer. The resulting $N_2$ gas was carried into the IRMS for isotopic ratio determination. Isotopic compositions are expressed in delta (δ) notation relative to internationally accepted atmospheric nitrogen (AIR) standards using the formula: $\delta^{15}N$ (‰) = $(R_{sample} - R_{standard}) / R_{standard} \times 1000$. Where $R_{sample}$ is the isotope ratio (i.e. $^{15}N/^{14}N$) of the sample, and $R_{standard}$ is that of the reference material. Variations in stable isotope ratios are reported as parts per thousand (‰) deviations from the AIR standard. The $N_2$ reference gas was calibrated using isotopic reference material (USGS40, L-glutamic acid, $\delta^{15}N=-4.52$ ‰) and was used as a reference gas for all measurements. The standard deviation of $N_2$ reference gas (n=10) was less than 0.06‰, ensuring high analytical precision (Supplementary Data 11). At the same time, 3-5 larvae from each group were sent to Scale Biomedicine Technology Co., LTD (Beijing, China), where the relative abundance of $^{15}N$-labeled amino acids was analyzed using UHPLC-HRMS/MS, the method was the same as described above (Supplementary Data 12).

All data were tested for normality using the Shapiro-Wilk *W*-test. Normally distributed data were analyzed using one-way ANOVA followed by Tukey's post-hoc tests, while non-normal data were assessed using Kruskal-Wallis tests and Dunn's Test for pairwise comparisons (see Supplementary Table 4). All statistical analyses were conducted in R version 4.1.1.

#### Measurement of Fitness Indicators

We evaluated colony fitness by measuring five key indicators. First, the survival rates of both workers and larvae were determined by counting the number of live individuals over time in the 30-day feeding experiment (Supplementary Data 11). The "coxph" function from the "survival" package v 3.4.0 in R was used to compare different treatment groups. Second, we monitored larval development by measuring larval body length daily using ImageJ v1.54g in the 30-day feeding experiment. To standardize the measurements, each daily body length value was divided by the mean body length recorded on Day 1 for the respective group (Supplementary Data 12). Generalized linear mixed models (GLMMs) were used to determine the fixed effects of the presence or absence of symbiotic bacteria in worker ants and development time on larval body length, with colony specified as a random effect. The models were constructed using the "glmer" function from the "lme4" package v1.1.35.5, and their significance was tested using ANOVA. In addition, we recorded pupation and eclosion rates (Supplementary Table 9) and also measured the head width of newly emerged workers using ImageJ v1.54g (the distance between the posterior regions of both eyes, a standard proxy for body size) as an indicator of body size (Supplementary Table 10) in the 90-day feeding experiment. Due to no statistically significant differences in the pupation and eclosion rates and worker head sizes between the $^{15}$N-urea-Sym-worker and $^{14}$N-urea-Sym-worker control groups (Supplementary Table 11), the two groups were subsequently combined into one group for statistical analysis in comparison with the $^{15}$N-urea-Apo-worker treatment group (Supplementary Table 3). Bartlett's test was used to assess homogeneity of variances. Normal data were compared using t-tests, and non-normal data were analyzed by Wilcoxon-Mann-Whitney test. All statistical analyses were conducted in R version 4.1.1.
